# Spatial and Single-cell Transcriptomics Reveal Programs Governing Fibroblastic Foci Fibroblasts

**DOI:** 10.1101/2025.09.26.678416

**Authors:** Yan Stein, Ophir Freund, Sarah Borsekofsky, Doron Cohn-Schwartz, Liran Levy, Efrat Ofek, Avishay Spitzer, Taylor Adams, Jonas C. Schupp, Naftali Kaminski, Amir Bar-Shai, Avraham Unterman

## Abstract

**Rationale:** Fibroblastic foci (FF) are central for fibrosis propagation in idiopathic pulmonary fibrosis (IPF), yet their resident fibroblast subpopulation remains poorly defined.

**Objective:** To deeply characterize FF-fibroblasts and identify programs governing their differentiation and activation.

**Methods:** GeoMx spatial transcriptomics was applied to IPF lung regions, comparing carefully delineated FF (excluding overlying epithelium) with areas of established fibrosis, to derive an FF-specific signature. This signature was projected onto a large single-cell RNA-sequencing (scRNA-seq) dataset to identify an FF-specific fibroblast cluster, facilitating its deep characterization.

**Measurements and Main Results:** GeoMx revealed 272 increased genes in FF; this FF-signature was validated against several independent spatial transcriptomics and proteomics datasets. FF-signature genes were highly enriched in a specific cluster of fibroblasts in the IPF scRNA-seq dataset. *MMP11* was the most specific gene of the FF-fibroblast cluster, and was validated using in situ RNA hybridization. Furthermore, higher MMP11 bronchoalveolar lavage protein levels were found in progressive pulmonary fibrosis and were associated with worse outcome in Cox regression (HR 4.54, 95% CI 1.24-15.8). Cell-cell interaction analysis with fibroblasts and adjacent aberrant basaloid cells, detected signals that may induce the FF-fibroblast signature, including from TGF-β superfamily ligands. Using trajectory and transcription factor (TF) analysis we show that the trajectory from non-FF to FF-specific fibroblasts is correlated with activation of EMT-associated TFs and deactivation of AP-1 family TFs.

**Conclusions:** We identified transcriptional programs governing FF-fibroblasts, for which *MMP11* serves as a novel highly-specific biomarker with prognostic significance. We highlight potential regulators whose targeting may curb pulmonary fibrosis progression.

## Introduction

Idiopathic pulmonary fibrosis (IPF) is characterized by excessive extracellular matrix (ECM) deposition, leading to progressive lung scarring with a median survival of 3-5 years (1). IPF lungs exhibit a histopathological pattern termed usual interstitial pneumonia (UIP) (2), characterized by unique histological structures termed fibroblastic foci (FF) (3), that are rich in active fibroblasts. FF are often found in the boundary between dense fibrotic tissue and histologically normal lung tissue (4). They are postulated to form the leading edge of fibrosis propagation in IPF (5), underscoring the need to elucidate the mechanisms driving their formation and function to inform new therapeutic strategies.

Currently, there is no good in vivo model for FF which makes it difficult to mechanistically study these structures. Despite the generation of multiple large single-cell RNA sequencing (scRNA-seq) datasets from lung tissues of dozens of pulmonary fibrosis (PF) patients (6–8), the subpopulation of fibroblasts that reside in FF has not been spatially resolved. Recent spatial transcriptomics studies have provided valuable insights about cell subpopulations and their spatial organization in PF (9–14), but lacked whole-transcriptome coverage and/or did not integrate spatially-resolved FF data with large-scale scRNA-seq datasets. Such integration should reveal the transcriptional programs governing FF-fibroblast differentiation and activation.

In this study, we utilized GeoMx digital spatial profiler (GeoMx DSP) (15) on IPF lung tissues, to characterize the transcriptional signature of the fibroblasts within FF, and validated it using independent spatial proteomics and transcriptomics datasets. Next, we used this signature to identify a cluster of FF-specific fibroblasts in a large scRNA-seq dataset derived from dozens of IPF patients (6). We identified *MMP11* as a novel highly specific marker of FF-fibroblasts and validated its specificity using in situ RNA hybridization (ISH). Furthermore, MMP11 protein levels in bronchoalveolar lavage (BAL) fluid were higher in patients with progressive pulmonary fibrosis (PPF) than in those without a progressive phenotype and were associated with worse outcome in Cox regression analysis, supporting their potential prognostic value. Cell-cell interaction analysis of FF-fibroblasts with fibroblasts and adjacent aberrant basaloid cells, detected signals that may induce the FF-fibroblast signature, including from TGF-β superfamily ligands. Using trajectory and transcription factor (TF) analysis we show that the trajectory from *MMP11*-negative to *MMP11*-positive (FF-specific) fibroblasts is correlated with activation of EMT TFs and deactivation of AP-1 family TFs. Taken together, our integrative spatial and single-cell approach was able to spatially-resolve and deeply characterize a rare FF-specific fibroblast subpopulation on a large scRNA-seq dataset. These data shed new light on the mechanisms that may lead to the propagation of PF, offer novel targets for therapeutic interventions, and suggest a novel FF-specific biomarker for fibrosis progression.

## Methods

### Participants

GeoMx spatial transcriptomics and ISH experiments were conducted on formalin-fixed and paraffin-embedded (FFPE) IPF/UIP lung samples previously obtained with informed consent on protocols approved by the Institutional Ethics Committees at Sheba (SMC-9101-22) and Tel Aviv Sourasky (16-660-TLV) Medical Centers. All patients (Table E1) had a UIP pattern as determined by expert pathologists (S.B. and E.O.) following international guidelines (16), with multiple FF. Diagnoses were established by multidisciplinary discussion in accordance with current guidelines (16, 17).

BAL was performed on patients with fibrotic interstitial lung disease (ILD) who underwent bronchoscopy as part of their ILD evaluation between 2021-2024 (Table E2). BAL collection, storage and downstream usage was approved by the Institutional Ethics Committee at Tel Aviv Sourasky Medical Center (21-0255-TLV), and all patients gave informed consent.

### Spatial profiling of IPF patient lung tissues

FFPE lung tissues of an IPF patient were profiled using GeoMx® DSP using Whole Transcriptome Atlas (WTA) probes, as previously described (15, 18). Slides were stained by hematoxylin and eosin (H&E) and were examined by a certified pathologist experienced in ILD (S.B.) to identify regions of FF and established fibrosis without FF, epithelial, or major vascular structures. The consecutive unstained tissue slices were sent to NanoString Technologies laboratories (Seattle, WA, USA) for further processing, regions of interest (ROI) capture, sequencing, and analysis.

### GeoMx data analysis

Data filtering and normalization were performed on GeoMx DSP Analysis Suite (V2.3.4). Differentially expressed gene analysis comparing FF to established fibrosis ROIs on log2-transformed normalized counts was performed using the R/Bioconductor package limma (19), to produce the FF-signature. Genes were considered increased if exhibiting log2FC >0.6 and p-value <0.05 adjusted for multiple comparisons using Benjamini-Hochberg (FDR) procedure.

### ScRNA-seq data analysis

A previous scRNA-seq dataset of IPF and control lungs by Adams et al. (6) was used for deep profiling of FF-fibroblasts, that were spatially-resolved using the FF-signature. Data was subset to include only cells originally annotated as myofibroblasts. Further downstream processing and analysis was performed using the R package Seurat, v4.0.4 (20). Differentially expressed genes were identified using ‘FindMarkers’ function, applying a log2FC cutoff >0.6 and adjusted p-value (FDR) <0.05. Pseudotime trajectory analysis was performed using ‘slingshot’ R package, and differentially expressed genes along the trajectory were inferred using ‘tradeSeq’ R package. TF analysis was performed using pySCENIC, and cell-cell interaction analyses were performed using ‘NicheNet’ and ‘CellChat’ algorithms.

### RNAscope ISH

ISH was performed using RNAscope 2.5 HD Reagent Kit Red and Hs-MMP11 (#479741) probe (Advanced Cell Diagnostics) according to manufacturer’s protocol.

### MMP11 BAL fluid ELISA

MMP11 ELISA was performed on 100 µl of the BAL supernatants using the Human MMP-11 ELISA colorimetric kit (NBP3-06887, Novus Biologicals) according to manufacturer’s instructions. Patients were divided to PPF group and a non-PPF group using the 2022 ATS/ERS/JRS/ALAT guidelines definition for PPF (17). Statistical analyses comparing MMP11 levels and survival analyses were performed by SPSS software v28 (IBM).

### Data availability

GeoMx raw data will be available in GEO upon publication. Normalized counts can be found in Table E3. Raw data of the re-analyzed scRNA-seq dataset (6) are available in GEO (GSE136831).

Additional details are found in the online supplementary materials.

## Results

### Spatial transcriptomics in IPF identifies a unique signature of FF fibroblasts compared to areas of established fibrosis

We hypothesized that by comparing the transcriptome of the active fibrotic tissue in FF to areas of less-active established fibrosis outside of FF, we will be able to identify the unique transcriptomic signature of FF-fibroblasts and use that to detect a subcluster of FF-fibroblasts in a large scRNA-seq dataset. To that end, we used IPF patient-derived lung slices and subjected them to the GeoMx DSP spatial transcriptomics method (15). For the selection of ROIs, IPF lung slices were stained by H&E as well as fluorescently stained for pan-cytokeratin, CD68, and alpha smooth muscle actin (α-SMA) (Figure 1A). Four carefully delineated FF ROIs (without their overlying epithelial layer) and 9 established fibrosis ROIs (without prominent FF, epithelial, or vascular structures) from our index IPF patient were analyzed using GeoMx DSP and principal component analysis (PCA) confirmed their separation into two groups (Figure 1B). A total of 1054 genes were differentially expressed between FF and established fibrosis areas (log2FC > 0.6, p < 0.05, Table E4), out of which 272 were increased in FF and 782 were increased in established fibrosis ROIs (Figure 1C-D). Many of the genes that were increased in FF have been previously associated with lung fibrosis, including ECM components such as various collagens (e.g. *COL1A1*) and periostin (*POSTN*) (Figure 1C-D and Figure E1A-B) (21, 22). Indeed, pathway enrichment analysis of genes increased in FF indicated enrichment of pathways pertaining to ECM deposition and organization (Figure 1E). Notably, *CTHRC1* which was previously found to be increased in FF (23), was increased in our data as well (Figure 1C-D and Figure E1C). Beyond genes already implicated in lung fibrosis and expressed in FF, we identified several previously unrecognized genes increased in FF, including ones that may play a role in the disease. These include *MMP11*, which will be subsequently discussed, and *HTRA1* (Figure 1C-D and Figure E1D-E). Although *HTRA1* was associated with fibrosis and TGF-β signaling in other organs (24, 25), as well as exhibiting prognostic value in IPF patients (26), it has not been mechanistically implicated in IPF pathology and to the best of our knowledge was not previously reported to be expressed in FF.

**Figure 1:**
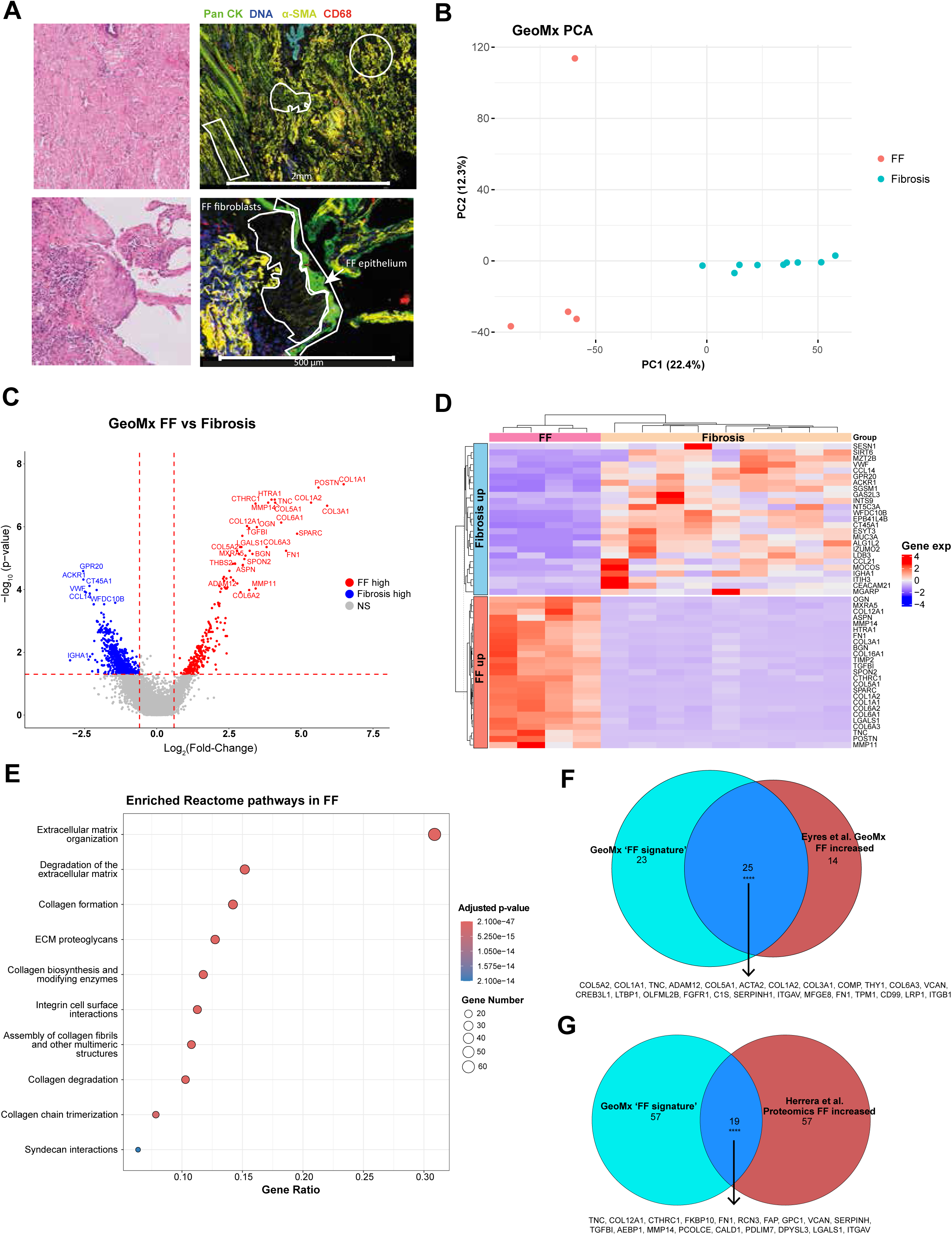
GeoMx DSP spatial transcriptomics identifies FF-increased genes. **(A)** Representative H&E and immunofluorescence images of ROIs from a lung of an IPF patient that were analyzed by GeoMx DSP. The ROIs that were analyzed are demarcated. The markers that were used for staining are as follows: Pan Cytokeratin (Pan CK, green), alpha-smooth muscle actin (α-SMA, yellow) and a nuclear stain (DNA, blue), and CD68 (red). **(B)** Principal component analysis (PCA) of gene expression from fibroblastic foci (FF) and established fibrosis ROIs. **(C)** A Volcano plot depicting statistically significant differentially expressed genes between fibroblastic foci and established fibrotic ROIs. Horizontal dashed line denotes p-value of 0.05, and vertical dashed lines denote log2FC of 0.6. **(D)** A heatmap showing the expression in the various ROIs of the top 25 genes upregulated in FF (‘FF up’) and top 25 genes upregulated in established fibrosis (‘Fibrosis up’) according to log2FC. Hierarchical clustering shows a clear separation between the ROIs. **(E)** Shown are the results of pathway enrichment analysis for genes increased in the FF ROIs among ‘Reactome’ pathway database. FF, fibroblastic focus; NS, not significant. (**F-G)** Shown are Euler diagrams depicting the intersections between our ‘FF signature’ and FF-increased genes (FDR < 0.05) that were reported by Eyres et al. spatial transcriptomics dataset (panel **F**), and between the ‘FF signature’ and FF-increased proteins (FDR < 0.05) reported by Herrera et al. spatial proteomics dataset (panel **G**). For both comparisons, **** denotes p<10^−4^ using permutation test for the observed Jaccard indices (0.4 for Eyres et al. comparison, 0.14 for Herrera et al. comparison).

We next sought to validate our spatial transcriptomics-derived FF-signature against independent spatial transcriptomics and proteomics datasets. We first compared it to two previous GeoMx spatial transcriptomics datasets by Eyres et al. (10) and Kim et al. (11). Eyers et al. compared FF to “adjacent [fibrotic] alveolar septae”, which is somewhat similar to our established fibrosis ROIs. This dataset included only a limited probe panel (1813 genes, out of which 1086 genes were detected), compared to our WTA panel (18,677 genes, 10,185 detected). To account for this difference, we only compared upregulated genes in FF in each study that were detected in both (n=795). Despite these differences, we observed a significant proportion of shared genes between our FF-signature and FF increased genes in the Eyers et al. dataset (Jaccard index = 0.4 vs. expected 0.03±0.02, permutation test p < 10^−4^, Figure 1F). Kim et al. used a WTA panel to compare FF to ‘fibrosis’ ROIs. Their FF increased genes exhibited a significant overlap with our FF-signature (Jaccard index = 0.29 vs. expected 0.005±0.003, p < 10^−4^, Figure E1F). In addition, we compared the FF-signature to an independent spatial *proteomics* cohort (27), which identified proteins that were increased in FF compared to areas of mature fibrosis outside FF, similar to our spatial design. Despite the use of a different modality (proteins vs mRNA), there was a significant overlap between the two datasets (Jaccard index = 0.14 vs. expected 0.003±0.004, p < 10^−4^, Figure 1G), which further corroborates the validity of our FF-signature.

To summarize, using GeoMx spatial transcriptomics we identified a set of genes increased in FF, termed the FF-signature, enriched for ECM deposition and organization pathways. This was achieved through a unique comparison of the highly active FF ROIs with established fibrosis ROIs, combined with whole-transcriptome coverage. We further validated this signature against independent spatial transcriptomics and proteomics datasets.

### The spatially-derived FF-signature identifies a specific subpopulation of fibroblasts within an IPF scRNA-seq dataset

Using the FF-signature, we wished to identify the fibroblast subpopulation that expresses this signature in a scRNA-seq dataset, in order to gain further insights based on a large cohort of patients. To this end, we utilized the scRNA-seq dataset of Adams et al. (6), derived from lung tissues of 32 IPF patients and 28 control subjects. We subclustered only the cells annotated as myofibroblasts from this dataset, because the FF-signature was recognized within this cell population, yielding 8 distinct clusters (Figure 2A). Differential gene expression analysis revealed that genes that were highly expressed in cluster 1 exhibited a significant overlap with the genes within the FF-signature (Fischer’s exact test, p < 2.2*10^−16^, Figure 2B-C, Figure E2 and Table E5). Furthermore, pathway enrichment analysis of cluster 1 genes identified similar pathways to those enriched in the FF-signature identified by GeoMx (Figure 2D). All in all, these data suggest that cluster 1 includes fibroblasts originating from FF.

**Figure 2:**
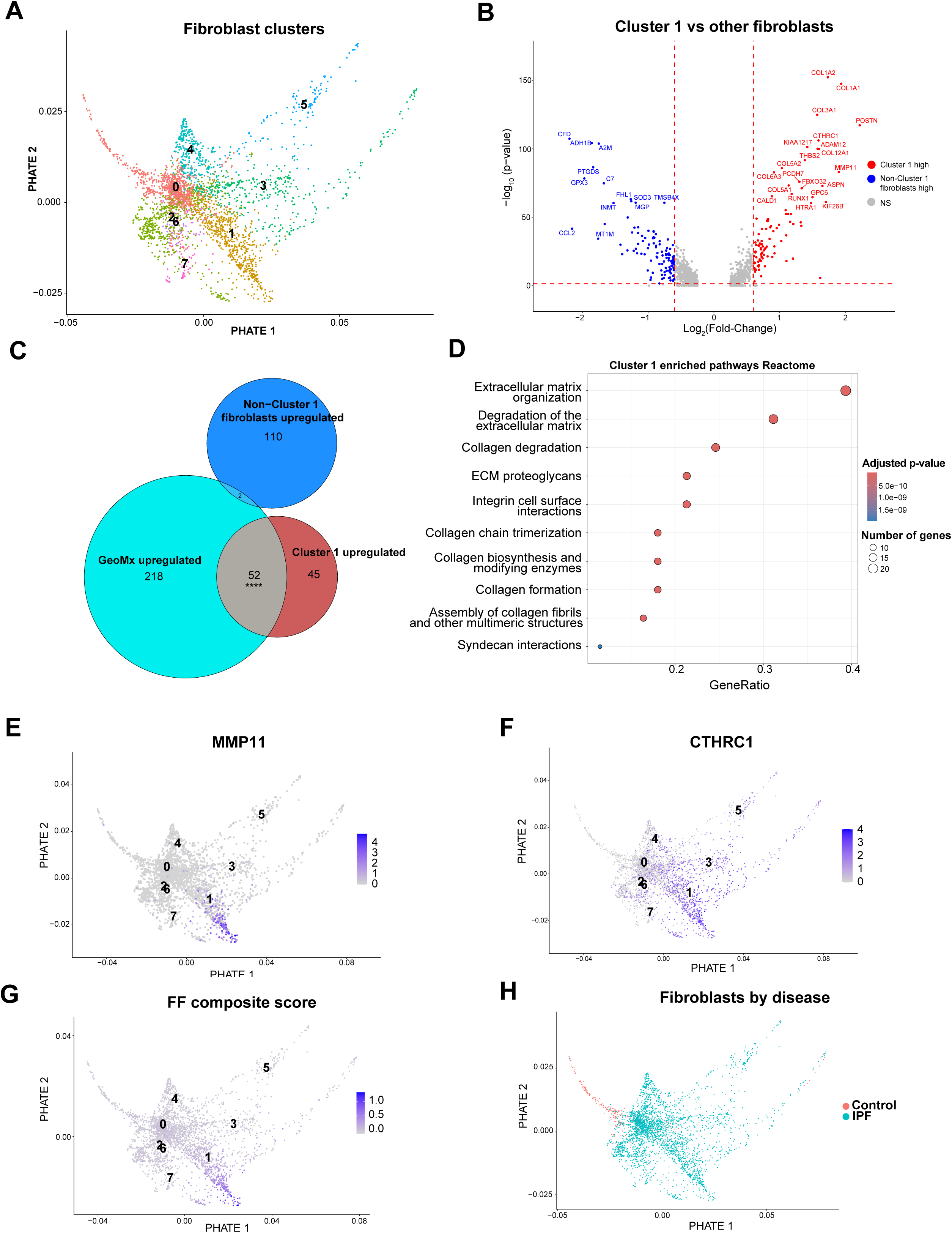
Genes increased in FF are expressed by a distinct subset of fibroblasts in IPF patients. **(A)** A PHATE representation of fibroblast subclustering data from Adams et al. IPF scRNA-seq dataset. **(B)** A volcano plot of differentially expressed genes between cluster 1 and other fibroblast clusters. **(C)** An Euler diagram showing the intersections of genes that were either upregulated cluster 1 (‘Cluster 1 upregulated’) or in the rest of clusters except cluster 1 (‘non-cluster 1 fibroblasts upregulated’) in and the genes that were upregulated in GeoMx FF (‘GeoMx FF upregulated’). **** denote p<2.2*10^−16^ (Fisher’s exact test) for the significant enrichment of cluster 1 upregulated genes among GeoMx FF upregulated genes. (D) Shown are the results of pathway enrichment analysis for genes increased in the cluster 1 among ‘Reactome’ pathways. Color signifies the significance level of the gene set enrichment, while dot size signifies the number of overlapping genes. **(E-F)** PHATE representations showing the expression patterns of *MMP11* (**E)** and *CTHRC1* (**F**) in the IPF fibroblast subclusters. **(G)** A PHATE representation of fibroblasts depicting the ‘FF composite score’, composed of the gene expression of the top 15 most specific genes to cluster 1. (**H)** PHATE representation of fibroblasts depicting the disease status (IPF or control) for each cell. Note that the majority of clusters, including cluster 1, are composed almost exclusively of cells derived from IPF patients.

To further refine the cluster to include specifically fibroblasts associated with FF, we calculated an ‘FF specificity score’ for each gene, based on the ratio of the average log2 fold-change between cluster 1 and the other clusters, and the proportion of cells that express the gene outside of cluster 1 (Table E5). This revealed a list of highly specific genes, such as *MMP11*, *MIAT* and *ENPP1*, which exhibited a highly specific expression pattern to a subset of cluster 1 (Figure 2E and Figure E3A-B). *CTHRC1*, a marker gene of high ECM-producing fibroblasts in a bleomycin murine PF model, is also expressed in FF in IPF lungs (23). While *CTHRC1* exhibited higher expression in cluster 1, it was also expressed in other fibroblast clusters (Figure 2F), and its specificity score was substantially lower than the three highly specific FF genes mentioned above. Next, we used the gene expression of the top 15 genes that had the highest ‘FF specificity score’ to compute a compound ‘FF score’ for each cell in the IPF fibroblast dataset. Interestingly, this revealed a subcluster within cluster 1 that was delineated by a high ‘FF score’ (Figure 2G and Figure E3C). We dubbed this subcluster of FF-fibroblasts ‘*MMP11+* cluster’ after *MMP11* that exhibited the highest ‘FF specificity score’, and focused on this subcluster in our subsequent analyses. Of note, this ‘*MMP11*+ cluster’ contains cells almost exclusively from IPF patients (335 cells vs only 4 cells in control subjects) (Figure 2H).

In summary, using the FF-signature derived from the spatial GeoMx data, we identified a cluster of fibroblasts in a scRNA-seq dataset of a large cohort of IPF patients that likely corresponds with FF fibroblasts. We further refined this cluster to delineate a highly specific subcluster of FF-fibroblasts, which is characterized by the expression of various FF specific genes, among which is *MMP11*.

### *MMP11* is a highly specific novel biomarker of FF-fibroblasts and its protein BAL levels are associated with worse prognosis

Our spatially-resolved scRNA-seq data suggests that *MMP11* is the most specific biomarker for the FF-fibroblast population. To further corroborate this finding, we performed ISH using RNAscope on lung samples of four patients with UIP (see Table E1 for patient baseline characteristics). Indeed, *MMP11* mRNA was found exclusively in FF in all four patients (Figure 3). Interestingly, in 2 out of 4 examined patients (#1 and #4, Figure 3), the expression pattern seemed to be limited to a subpopulation inside the FF that was in close proximity with the epithelial layer of the FF, which is populated by the previously characterized aberrant basaloid cells (ABCs) (6). Of note, based on the scRNA-seq dataset (6), *MMP11* is highly specific to FF-fibroblasts across all types of lung cells, including epithelial, immune, and stromal cells (Figure E3D). Using a human lung fibroblast cell-line, we show that TGF-β, a master fibrosis regulator (28), not only induces classical fibroblast activation genes such as *ACTA2* (α-SMA) and *COL1A1* (Figure E4A-B), but additionally upregulates *MMP11* (Figure E4C) at two different time-points.

**Figure 3:**
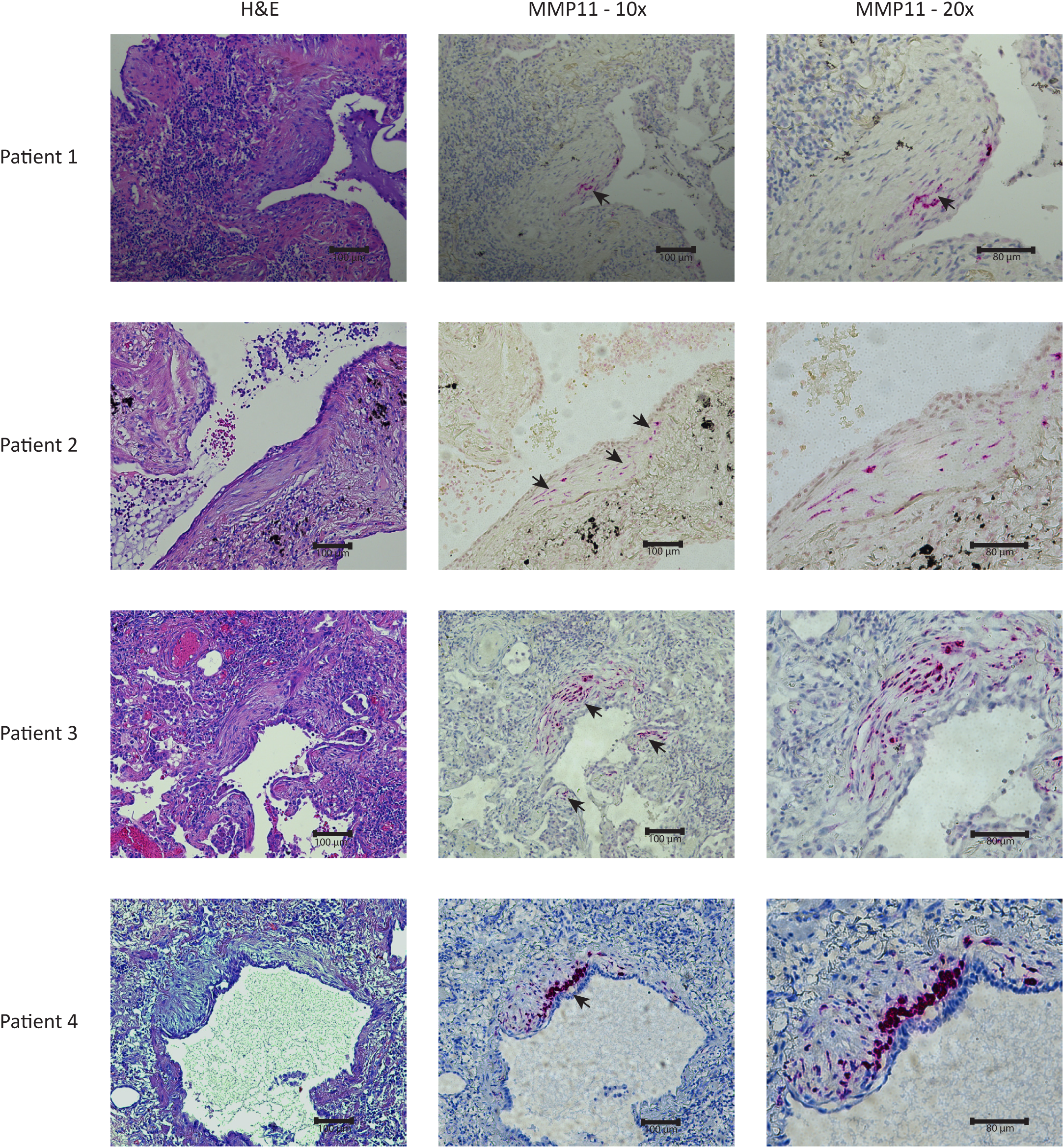
*MMP11* is a specific marker for FF subpopulation. *MMP11* mRNA was probed using RNAscope in-situ hybridization in sections from 4 different patients exhibiting UIP histological pattern. Shown are representative images of *MMP11* staining in 10x (middle panel) and 20x (right panel) magnifications, as well as an H&E staining of a consecutive slide (left panel). Arrowheads point to positive-stained cells.

Since the number and area of FF in PF is correlated with prognosis (29), we speculated that MMP11 protein levels will be higher in BAL fluid of fibrotic ILD patients with a PPF phenotype compared to those without progressive disease. We tested this hypothesis on BAL fluid samples of 33 fibrotic ILD patients (mean age 65.9 ± 13.1 years, 73% males, 52% with PPF; See Table E2 for baseline and clinical characteristics). Indeed, MMP11 protein levels were increased in PPF (Figure 4A), suggesting a possible prognostic role for this biomarker. Interestingly, the highest levels were detected in an IPF patient that was sampled during an acute exacerbation.

**Figure 4:**
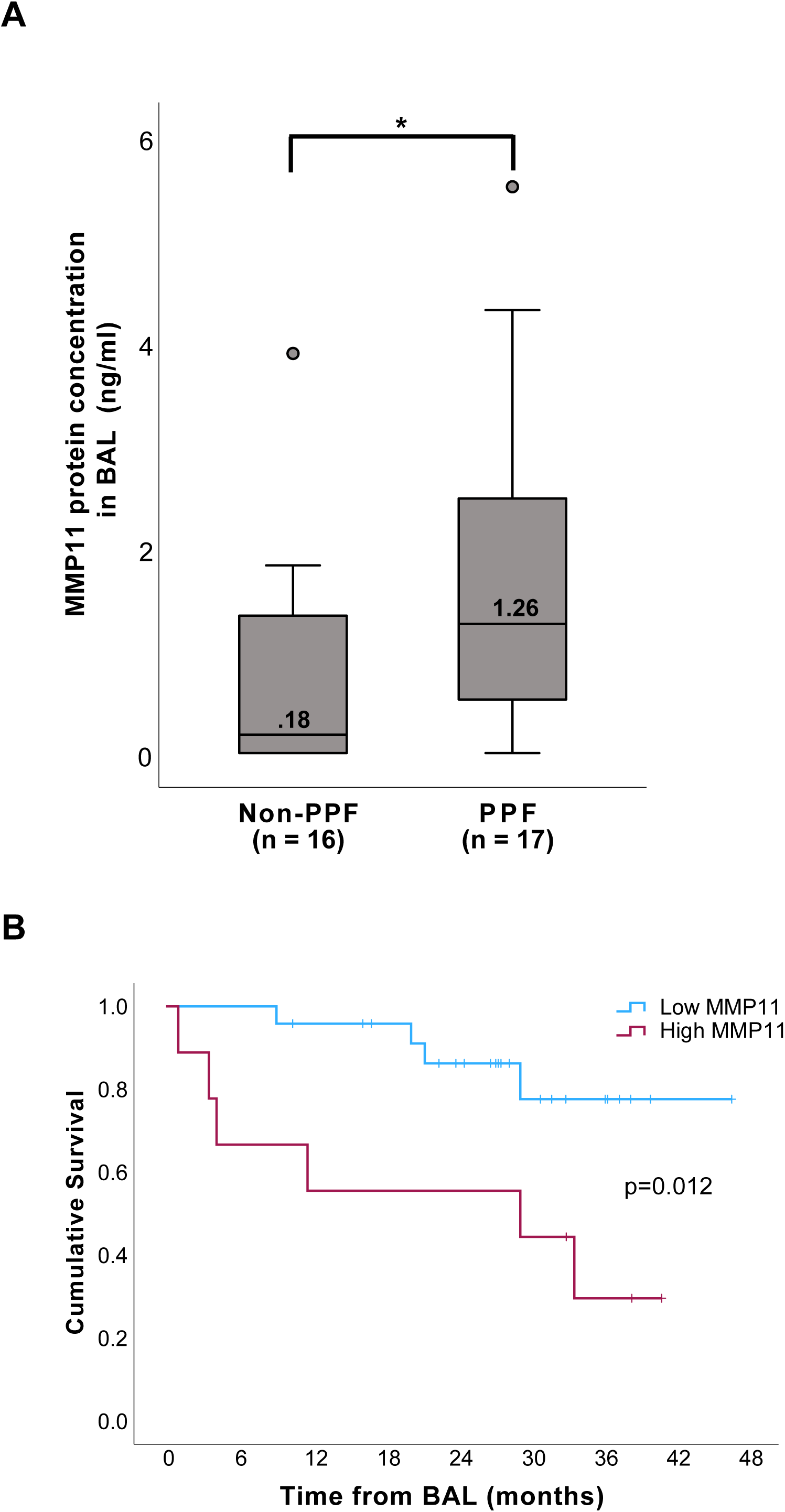
MMP11 protein levels are increased in bronchoalveolar lavage (BAL) fluid of patients with progressive pulmonary fibrosis (PPF), and high MMP11 levels correlate with poorer clinical outcome. (**A)** MMP11 protein levels were measured in BAL fluid of pulmonary fibrosis patients using ELISA. Shown are MMP11 levels in PPF patients compared to non-PPF patients. * denotes p<0.05 (Mann-Whitney U test). (**B)** A Kaplan-Meier curve depicting adverse event-free survival (defined as absence of transplantation, death or hospitalization due to exacerbation) of MMP11 high and MMP11 low patients, with group cutoff set at 1.6 ng/ml. The denoted p-value was calculated using the log-rank test.

To further validate MMP11 protein levels as a prognostic biomarker, we analyzed the association between BAL MMP11 protein levels and the combined outcome of death, lung transplantation, or hospitalization with ILD exacerbation using Cox regression. Higher levels of MMP11 as a continuous variable were associated with higher hazards for mortality (for every increase of 1 ng/ml in MMP11 levels, HR 1.46, 95% CI 1.09-1.96). We then divided the patients to high and low MMP11, using the cutoff of 1.6 ng/ml. Patients with high MMP11 had significantly higher hazard for the combined outcome (Figure 4B, HR 4.54, 95% CI 1.24-15.8).

In summary, our data indicates that MMP11 is a highly specific novel biomarker of a fibroblast subpopulation within FF, and that its protein levels in BAL fluid may be associated with progressive disease and worse clinical outcome.

### ABCs, fibroblasts and myofibroblasts exhibit cell-cell interactions with *MMP11*+ fibroblasts that may induce the FF-signature

In order to examine potential biological mechanisms leading to the emergence of the *MMP11*+ fibroblasts, we examined their cell-cell interactions. As these fibroblasts reside in the FF, we focused our search on cell types that are found in close proximity to these cells, which are other fibroblasts, myofibroblasts, and ABCs. First, we used NicheNet algorithm (30) to predict the potential ligands that are expressed in the aforementioned cell types and may induce the FF-signature. As expected, at the top of these ligands was TGF-β1, a well-known master regulator of lung fibrosis (28). Other ligands of the TGF-β superfamily, such as TGF-β2 and TGF-β3, BMP4 and BMP2, *INHBA* (encoding a subunit of Activin and Inhibin) were also predicted to induce at least part of the FF-signature genes (Figure 5A-C). In addition, IGF2 and FGF family ligands, which were previously implicated in PF (31, 32), were also predicted as potential ligands that induce the FF-signature (Figure 5A-C).

**Figure 5:**
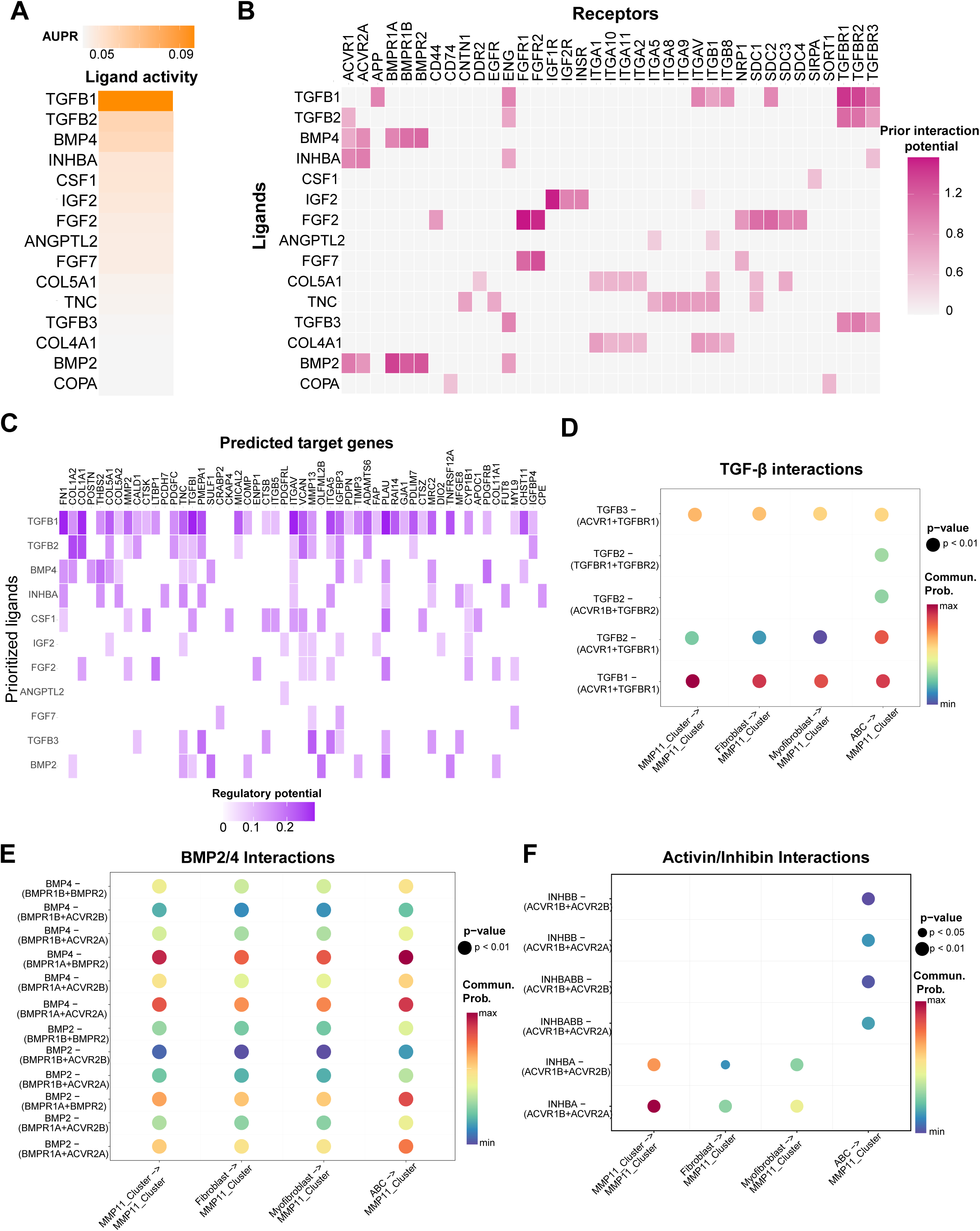
Cell-cell interaction analyses of *MMP11*+ fibroblasts. **(A)** Top ligands predicted to induce the ‘FF signature’ by ‘NicheNet’ algorithm. AUPR, Area Under the Precision-Recall Curve. **(B)** Ligand-receptor interactions that were predicted to induce the ‘FF signature. **(C)** Shown is the regulatory potential of various genes from the ‘FF signature’ by the top predicted ligands. **(D-F)** Specific ligand-receptor interactions in *MMP11*+ fibroblasts (‘*MMP11*+ Cluster’), non-*MMP11*+ Fibroblasts (‘Fibroblasts’), fibroblasts and ABCs that were predicted by ‘NicheNet’ were validated by ‘CellChat’ algorithm. Shown are interactions of ligands from the TGF-β (**D**), BMP2/4 (**E**) and Activin/Inhibin (**F**) families. Bubble color corresponds to communication probability (interaction strength) and bubble size corresponds to interaction’s p-value.

In order to validate these findings using another computational pipeline, and to examine which cells exhibit this interaction, we examined the cell-cell interactions between the *MMP11*+ fibroblasts and the aforementioned cells using ‘CellChat’ (33). Indeed, most of the major interactions that were predicted to induce the FF-signature were detected in fibroblasts, myofibroblasts and ABCs (Figure 5D-F, Figure E5A-C). Interestingly, some signals were strongly detected in an autocrine loop between the *MMP11* fibroblasts and themselves, indicating a potential maintenance mechanism. In summary, we were able to detect cell-cell interactions between *MMP11* fibroblasts and other neighboring cells, including ABCs, which may partially induce the FF-signature.

### Trajectory analysis from non-FF to FF fibroblasts correlates with activation of EMT-associated TFs and deactivation of AP-1 family TFs

Next, we performed a pseudotime analysis, in order to identify a potential trajectory by which non-*MMP11*+ fibroblasts may differentiate into *MMP11*+ fibroblasts, or vice versa. We identified a disease-related trajectory from fibroblast subcluster #4, which terminated in fibroblast cluster 1 and the ‘*MMP11+* cluster’, further denoted as ‘FF trajectory’ (Figure 6A top panel). We next examined the association of FF-signature genes with the ‘FF trajectory’ (Figure 6A bottom panel, Table E6). *LITAF*, which was shown to negatively regulate cardiac hypertrophy and fibrosis in mice (6), was decreased along this trajectory (Figure 6A bottom panel). As expected, genes encoding ECM proteins that were identified as part of the FF-signature, such *FN1*, *POSTN, CTHRC1* and various collagens, exhibited an increasing expression pattern throughout the trajectory (Figure 6A bottom panel). *MMP11* was only expressed at the cells in the final tip of the ‘FF trajectory’ (Figure 6A bottom panel).

**Figure 6:**
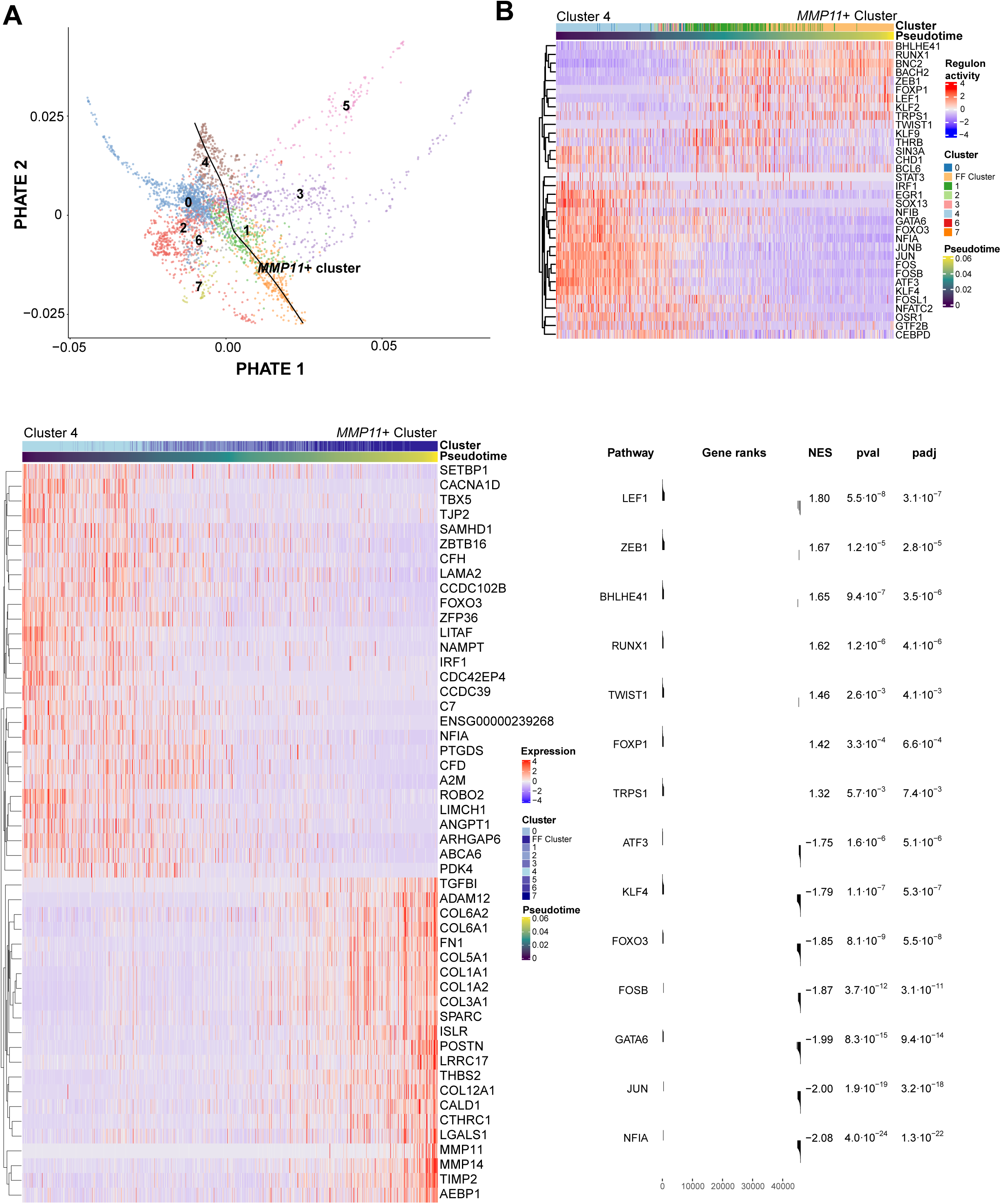
Transcription factor inference of genes associated with the transition between non-FF fibroblasts to MMP11-positive fibroblasts. **(A)** Top panel: A PHATE representation showing an inferred trajectory between non-FF fibroblasts and the ‘*MMP11+* cluster’. Bottom panel: A heatmap depicting the top 50 genes that exhibit the highest correlation with the ‘*MMP11*+ cluster’ trajectory. **(B)** Top panel: A heatmap exhibiting the inferred activity of transcription factors regulating the genes of the ‘*MMP11*+ cluster’ trajectory. Bottom panel: Shown are the results of a gene-set enrichment analysis (GSEA) that was performed for the enrichment of the inferred transcription factor target genes (regulons) among differentially expressed genes of the ‘*MMP11+* cluster’. NES, normalized enrichment score; pval, p-value; padj, adjusted p-value (FDR).

To further explore which upstream regulators were associated with the progression of the trajectory and the emergence of the *MMP11+* fibroblasts, we performed a transcription factor (TF) inference for genes that were either significantly associated with the ‘FF trajectory’ or were differentially expressed between the beginning and the end of the trajectory. This yielded TFs and their potential target genes, hereby referred as regulons, which were positively correlated with the ‘FF trajectory’, as well as negatively correlated TFs (Figure 6B top panel and Table E7). Gene-set enrichment analysis (GSEA) indicated a high enrichment of the TF regulons among the FF-signature genes (Figure 6B bottom panel), and there was a significant overlap between the predicted regulons and the FF-signature (Figure E6).

Strikingly, many of the TFs that were positively associated with the ‘FF trajectory’ are epithelial-to-mesenchymal transition (EMT) TFs, including *TWIST1* and *ZEB1,* which are also implicated in fibroblast activation and fibrosis (34, 35). Furthermore, *TWIST1* itself was upregulated in cluster 1, and ranked 10^th^ in its specificity score (Figure E2 and Table E5). In contrast, TFs negatively associated with the ‘FF trajectory’ were enriched with activator protein-1 (AP-1) family members, such as *JUN, JUNB, FOS, FOSB*, *FOSL1* and *ATF3*. Notably, *NFIA,* which has a regulon that exhibited highly significant negative association with ‘FF trajectory’, also exhibited significant decreased gene expression of the TF itself (Figure 6A bottom panel). The implications of these findings will be elaborated in the discussion section.

## Discussion

Fibroblastic Foci have been postulated to form the leading edge of fibrosis propagation in UIP, instigating an interest in the biological mechanisms underlying their formation and activation. Despite the existence of several large scRNA-seq datasets encompassing dozens of patients (6–8) and spatial transcriptomics datasets (9–14), none of them was able to extensively characterize the activated fibroblast subpopulation residing in FF. In the current work, we undertook an approach involving both spatial transcriptomics and scRNA-seq that facilitated the identification and deep characterization of FF-fibroblasts, and suggest potential regulators that may be responsible to their formation and activation.

*MMP11* was identified as the most specific transcript to the FF-fibroblast subpopulation, obtaining higher ‘FF specificity score’ than previous FF biomarkers like *CTHRC1*. Using ISH we show that-at least in some patients-it preferentially localizes to fibroblasts that are in close proximity to ABCs. *MMP11* belongs to the matrix metalloproteinase (MMP) family of proteins, and unlike most other MMPs, which are secreted as inactive zymogens, it is cleaved intracellularly and secreted in its already active form (36). It has been well established that the number and area of FF in biopsy correlate with prognosis (29), yet a minimally-invasive and specific FF-biomarker was not previously available. Our current work suggests that MMP11 BAL fluid level may serve as a clinically-relevant prognostic biomarker in PF, since it was elevated in PPF and was outcome-predictive using Cox regression and Kaplan-Meier analyses.

Several previous studies have attempted to perform spatial transcriptomics on FF. One such study (22) used laser capture microdissection in conjunction with bulk RNA-seq to characterize FF in IPF patients. In this study, the authors did not compare FF to another ROI, but rather employed a bioinformatics approach to identify transcriptional programs that correlate with the expression of collagen-encoding genes. This led to identification of several pathways, such as the TSC2/RHEB, as activated in FF. However, because this analysis lacked a comparator ROI, it likely overlooked transcriptional programs not directly correlated with collagen expression. More recently, there have been several studies (9–14) employing spatial transcriptomics in IPF patients, examining, among others, FF ROIs. However, there are two notable distinctions between the current study and those studies. First, as these studies performed multiple comparisons among many ROIs, they did not perform an elaborate comparison specifically between FF and established fibrotic areas. For example, Blumhagen et al. (9) compared ‘transition zones’, which include FF but also other epithelial regions, with other ‘normal looking parenchyma’. Others, such as Eyres et al. (10), compared FF regions with adjacent regions but not specifically with established fibrosis areas, and used a small panel of probes limiting the transcriptional characterization of these ROIs. Our focus on the comparison of FF and dense fibrosis facilitated the identification of genes and transcriptional programs that are specifically associated with fibrosis initiation and propagation, rather than fibrosis maintenance. The second difference is that our spatial transcriptomics analysis was complemented by the identification and elaborate characterization of *MMP11*+ FF-fibroblasts using scRNA-seq data, including their interactions with neighboring cells, and potential programs involved in the transition of fibroblasts to FF-fibroblasts, or vice versa. While Franzen et al. (12) mention *MMP11*, among other ECM genes, as increased around regions that include ABCs, we provide the first evidence for MMP11 as a specific FF-fibroblast biomarker and highlight its potential prognostic relevance in PF.

Our cell-cell interaction analysis identified a complex network of ligands that were previously implicated in PF, such as TGF-β ligands, Activin/Inhibin (encoded by *INHBA*), BMP2 and BMP4 as well as proteins from the IGF and FGF families. In the context of PF, it was observed that BMPs typically repress ECM production, and that its levels are attenuated in PF in a bleomycin model (37); however, BMP signaling is highly complex and context-dependent (38). In contrast, Activin A (composed of two subunits encoded by *INHBA*), which according to our data seems to exert its activity via ACVR1B an ACVR2A, is generally known to be pro-fibrotic in liver, kidney and muscle (39). Intriguingly, Activin A expression was previously observed in epithelial cells in the vicinity of FF, likely representing the ABCs (40). Furthermore, Activin A levels were increased in the plasma of IPF patients with acute exacerbation compared to IPF patients with stable disease (41). In our data, the strongest interaction of *INHBA* with *ACVR1B* and *ACVR2A* was observed as autocrine loop between *MMP11+* fibroblasts and themselves, indicating a potential positive feedback loop maintaining the enhanced pro-fibrotic activity in FF.

Our study identified potential TFs that may regulate the transition between non-FF fibroblasts and *MMP11+* FF-fibroblasts. Interestingly, this transition was accompanied by an increased activity of TFs associated with EMT. ABCs, that reside in close proximity to FF, were reported to activate EMT programs which may indicate a shared signaling activating these programs in both cell types. Some TF associated with EMT, such as *ZEB1* (*42*) and *BHLHE40/DEC1* (43), were previously implicated in promoting PF in bleomycin mouse model. Interestingly, chromatin accessibility study indicated that IPF myofibroblasts exhibited significantly more open chromatin in *TWIST1* promoter than IPF and control fibroblasts, which also resulted in an increase in its expression (44). In addition, this study also reported that *Twist1* overexpression in *Col1A2* fibroblasts in a bleomycin mouse model resulted in increased collagen synthesis, suggesting a role for *TWIST1* in fibroblast activation (44). Our data is in agreement with these results, and indicates that more specifically, *TWIST1* and other EMT TFs may be associated with the emergence of FF and fibrosis propagation. In contrast to EMT TFs, AP-1 family of TFs exhibit a negative correlation with the trajectory of *MMP11+* FF-fibroblasts, including *JUN*, *JUNB*, *FOS*, *FOSB*, *FOSL1* and *ATF3*. *FOSL1* (also known as *FRA1*) was shown to negatively regulate PF in a bleomycin model in vivo (45), while other reports regarded AP-1 TFs as profibrotic (46–48).

Our study is not without limitations. First, our FF-signature is based on a limited number of ROIs. We mitigate this with multiple external validations using spatial transcriptomics and proteomics datasets. Second, since FF are absent in pre-clinical models of PF, we could not validate our findings in vivo. Lastly, developing MMP11 as a clinically-ready biomarker was beyond our scope; future prospective studies should standardize its measurement, and validate its prognostic utility.

To conclude, our study identified an FF-signature and deeply characterized the fibroblast population residing in FF. We identified multiple cell-cell interactions and TFs that may regulate their differentiation, and highlight *MMP11* as a specific marker of FF-fibroblasts and potentially as a prognostic biomarker of fibrosis progression and outcome. Potential future directions may involve the generation of FF-like fibroblasts in vitro and their co-culture with epithelial cells or ABC-like cells using 3D models such as organoids. Furthermore, genetic ablation or pharmaceutical perturbations of TFs or cell-cell interactions may be performed on precision-cut lung slices (PCLS). Such mechanistic validations may pave the way towards the development of therapeutic interventions that block FF formation and fibrosis propagation.

## Acknowledgements

We would like to thank Aaron McClelland, Stefan Rogers and the Technology Access Program (TAP) team at NanoString Technologies (Seattle, WA, USA) for performing sample processing and raw data analysis of GeoMx spatial transcriptomics data.

## Author contributions

Y.S. and A.U. conceived and designed the work. Y.S., S.B. and A.U. were involved in the GeoMx spatial transcriptomics sample preparation and analysis. Y.S, S.B. and E.O. contributed to sample preparation and performance of in situ RNA hybridization. Y.S., O.F., T.A., J.C.S, A.S and A.U. performed computational data analysis. Y.S., O.F. and D.C.S. performed experimental data analysis. L.L., A.B.S. and A.U. provided patient samples and relevant clinical data. N.K. provided single-cell RNA sequencing data. Y.S, O.F. and A.U. drafted the initial manuscript. All authors contributed to drafting the manuscript or revising it critically for important intellectual content and approved the final draft. A.U. supervised the project.

---

**Figure E1.**
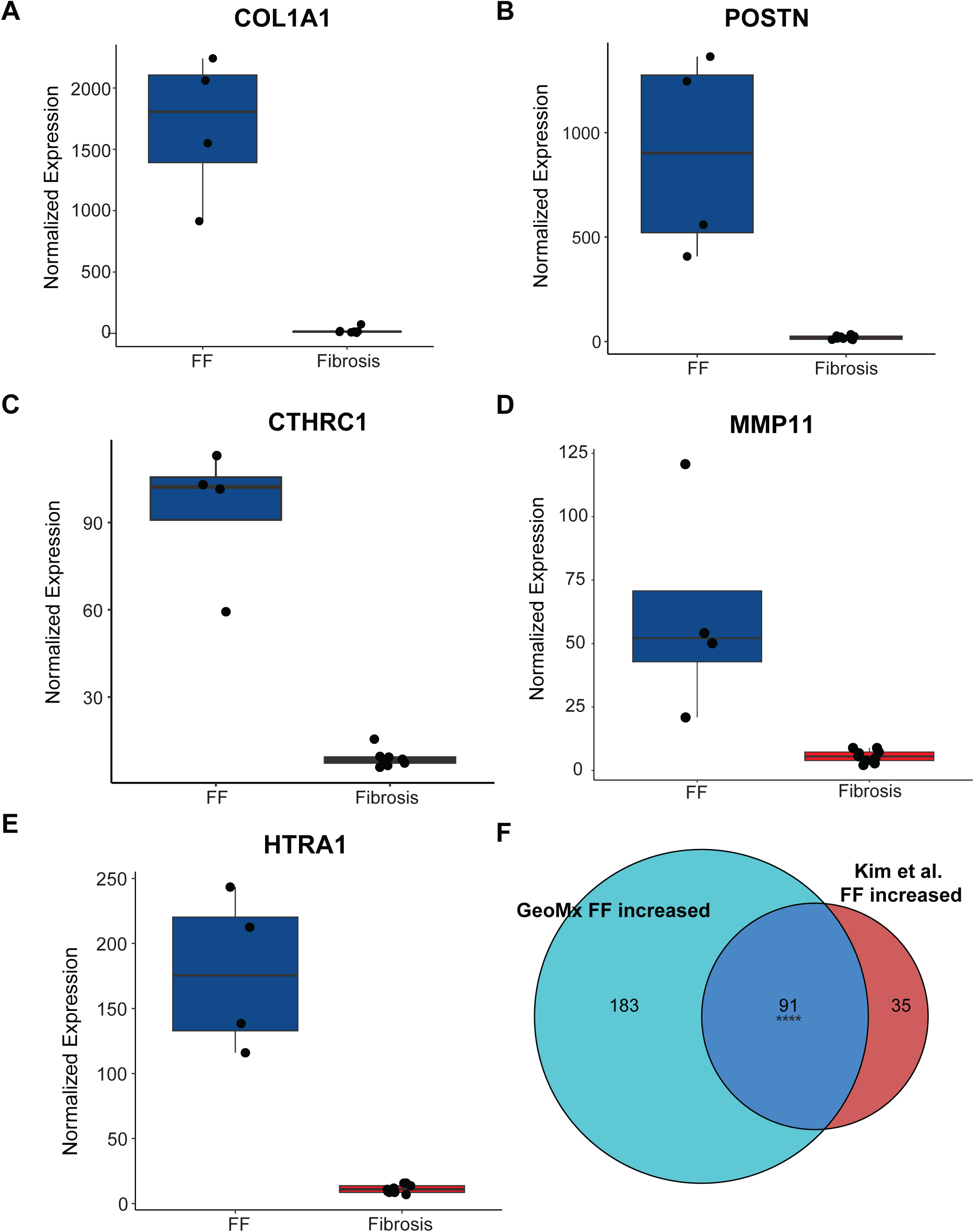
Various genes are increased in FF and exhibit a significant overlap with previously reported FF-increased genes in a spatial transcriptomics dataset. (**A-E**) Boxplots depicting the gene expression of various genes increased in FF vs established fibrosis (fibrosis) ROIs. Each dot represents an ROI. (**F**) An Euler diagram depicting the overlap between the genes that were increased in FF in our data and FF-increased genes from a spatial transcriptomics dataset reported by Kim et al. **** denote p < 10^−4^ (permutation test) for the overlap.

**Figure E2.**
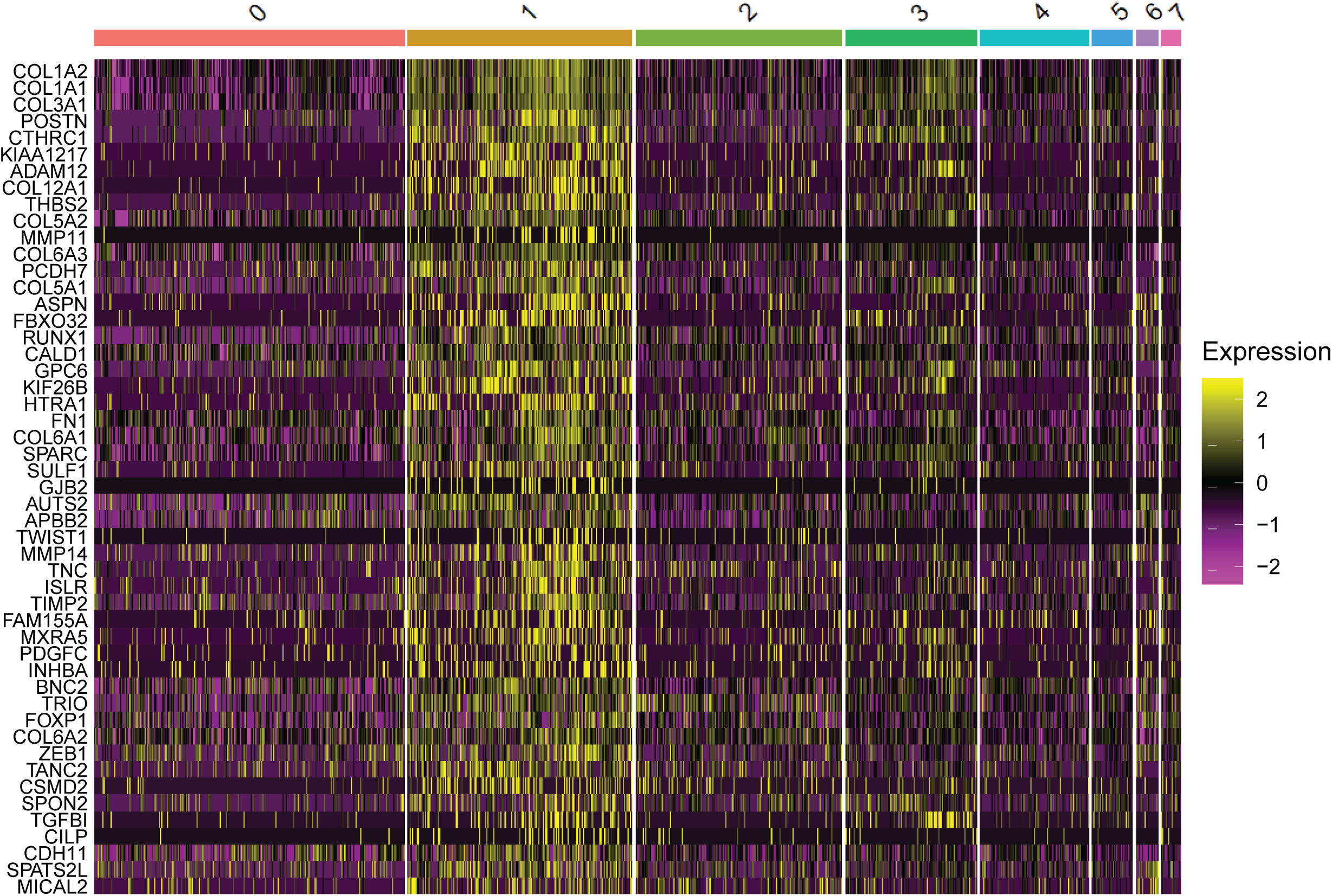
A heatmap of top 50 genes increased genes in fibroblast cluster #1. Genes are sorted according to adjusted p-value (Bonferroni correction), from the lowest (most significant) to the highest (least significant).

**Figure E3.**
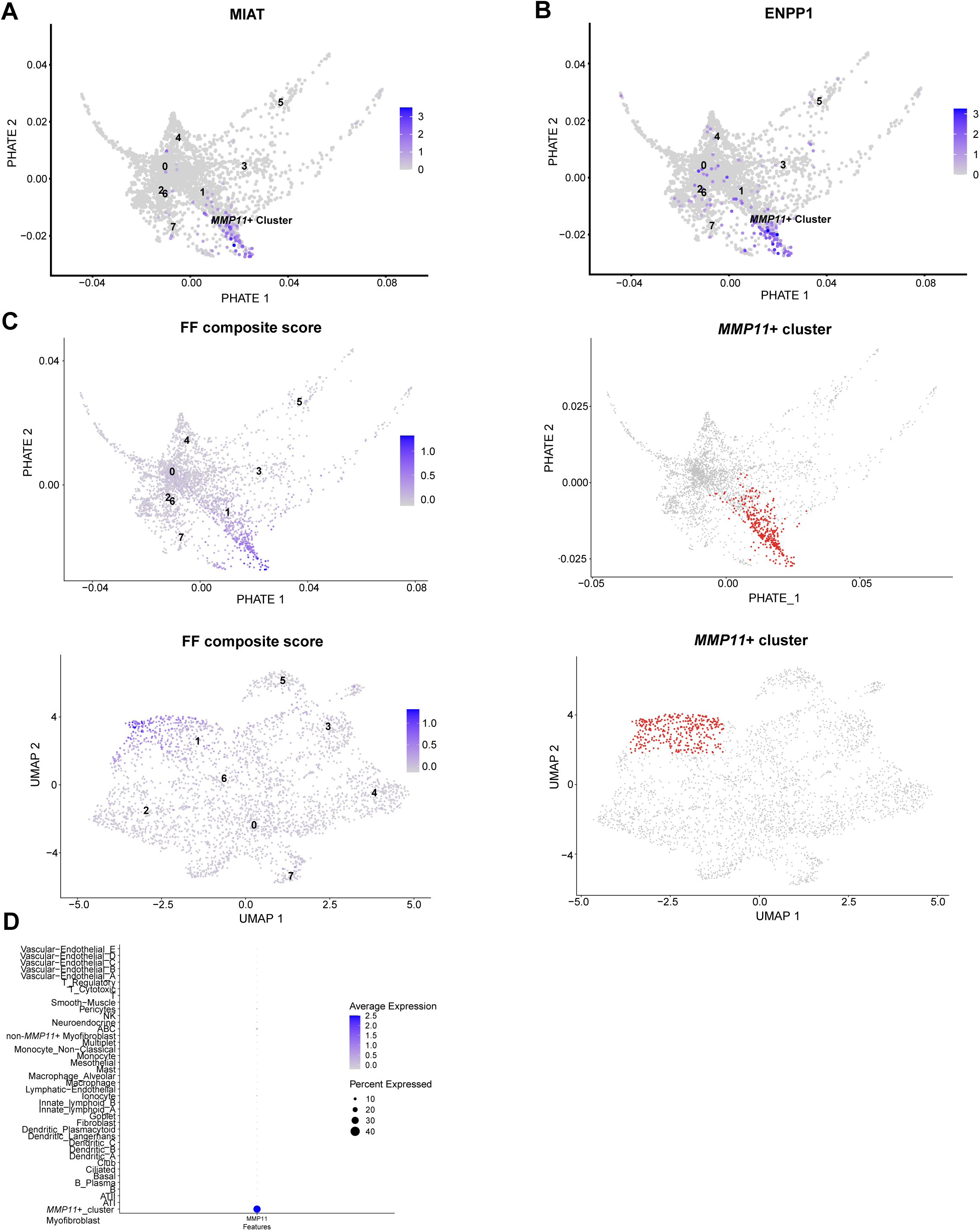
Delineation and characterization of *MMP11*+ cluster. **(A-B)** UMAPs showing the expression pattern of *MIAT* **(A)** and *ENPP1* **(B)** in myofibroblast clusters. **(C)** PHATE (top) and UMAP (bottom) plots showing FF composite score (left) and marked *MMP11*+ cluster (right) that was demarcated based on this score. **(D)** A dotplot showing the expression of *MMP11* along the whole cell types identified in the IPF cell atlas by Adams et al. The myofibroblast cluster was separated into non-*MMP11*+ myofibroblasts and *MMP11*+ cluster myofibroblasts. ABC, aberrant basaloid cells; NK, natural killer cells.

**Figure E4.**
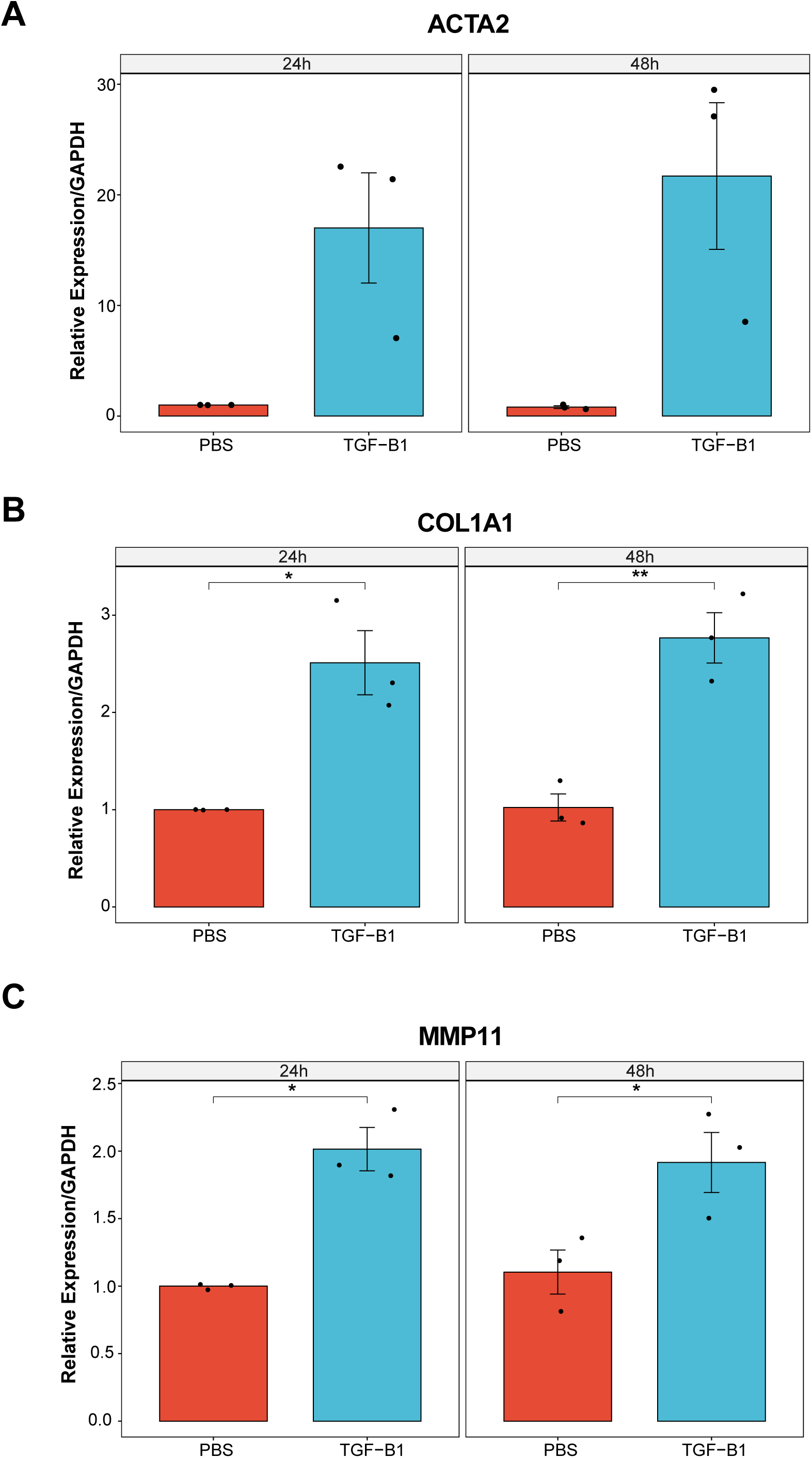
*MMP11* is upregulated by TGF-B1. WI-38 cells were treated with TGF-B1 (10 ng/ml) or PBS vehicle control. Cells were harvested at the specified time points, and the expression of the indicated genes was assessed by RT-qPCR. Gene expression was normalized to *GAPDH* housekeeping gene and to the corresponding 24h PBS treated sample. Individual data points indicate independent replicates (n=3), and error bars denote standard error of the mean (SEM). * denote p < 0.05 and ** denote p < 0.01 as determined by t-test.

**Figure E5.**
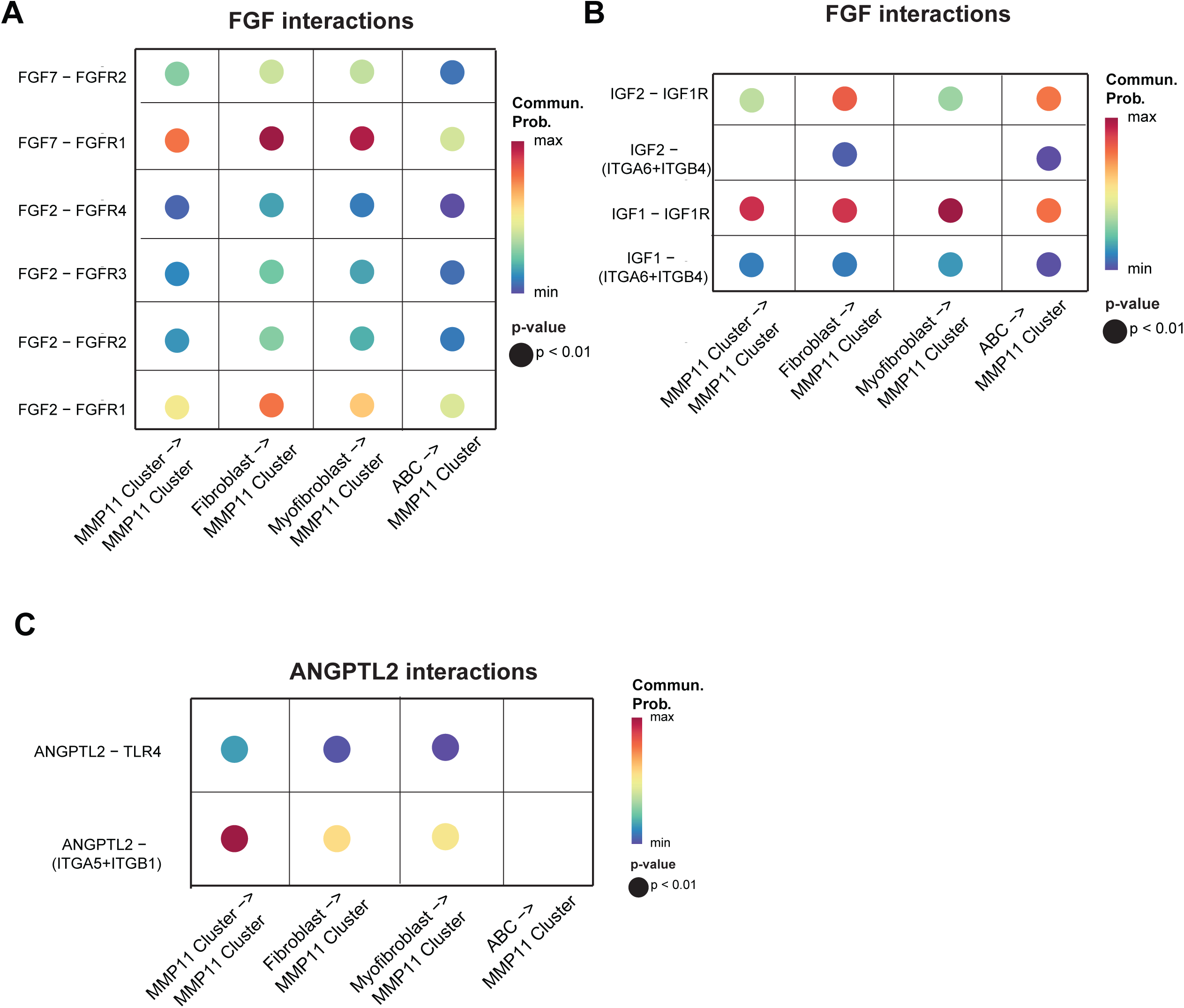
Cell-cell interactions inferred by ‘CellChat’. Shown are bubble plots of interactions of ligands from the FGF (A) and IGF (B) families, as well as ANGPTL2 interactions (C). Bubble color corresponds to communication probability (interaction strength) and bubble size corresponds to interaction’s p-value. Cell type annotation was borrowed from the original annotation by Adams et al. IPF scRNA-seq dataset. ABC, aberrant basaloid cells.

**Figure E6.**
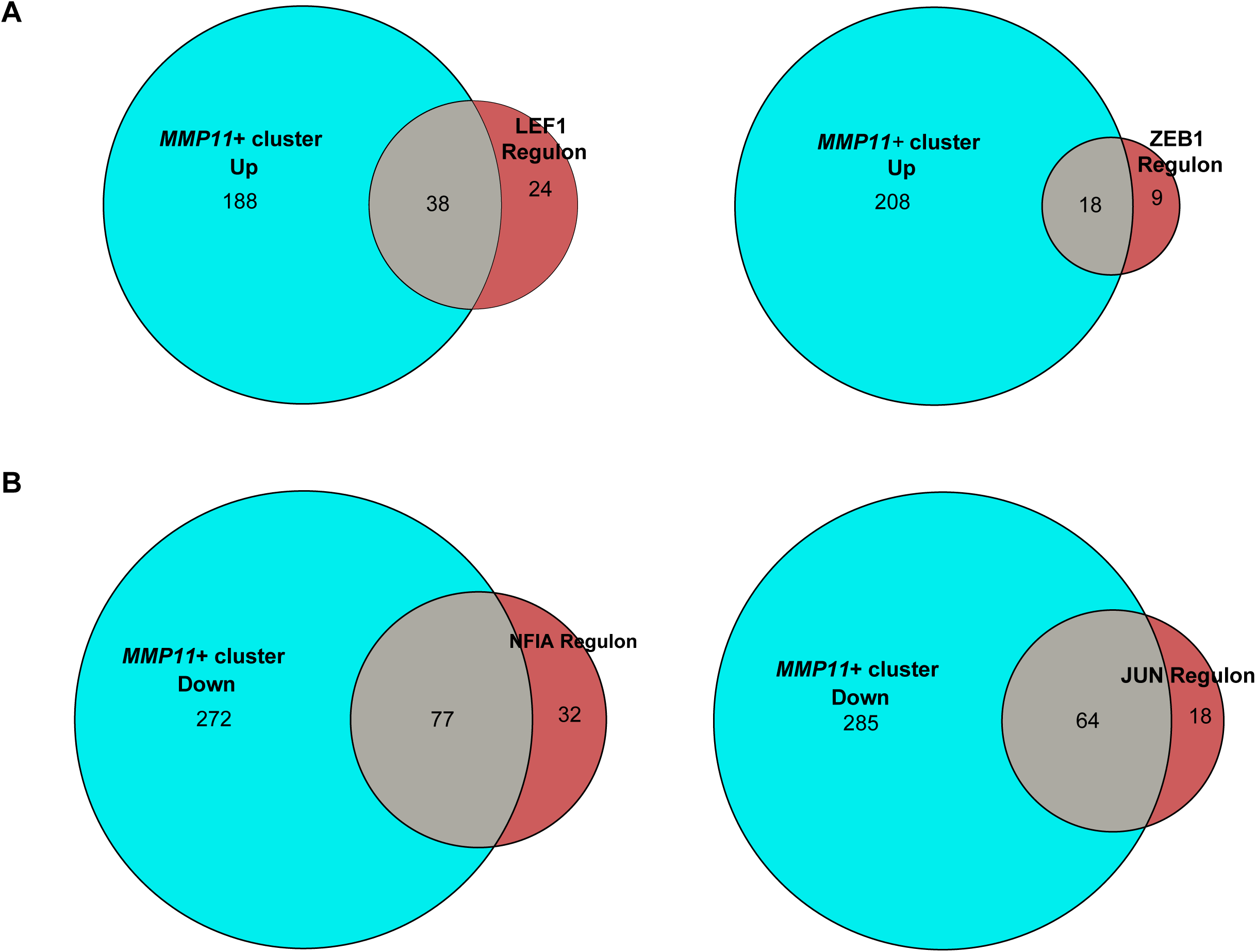
Enrichment of various transcription factor regulons among *MMP11*+ cluster differentially expressed genes. **(A)** Venn diagrams showing the intersection between genes that were increased in the *MMP11*+ fibroblast cluster (‘*MMP11*+ cluster up’) compared to other fibroblast clusters and TF regulons that were positively correlated with the ‘FF trajectory’. **(B)** Venn diagrams showing the intersection between genes that were decreased in the *MMP11*+ fibroblast cluster (‘*MMP11*+ cluster down’) compared to other fibroblast clusters and TF regulons that were negatively correlated with the ‘FF trajectory’.

