## Supplementary Methods for "Spatial and Single-cell Transcriptomics Reveal Programs Governing Fibroblastic Foci Fibroblasts"

**Participants and procedures**

GeoMx spatial transcriptomics and in situ RNA hybridization experiments were conducted on formalin fixed and paraffin embedded pulmonary fibrosis lung samples previously obtained with informed consent on protocols approved by the Institutional Ethics Committees at Sheba Medical Center (SMC-9101-22) and Tel Aviv Sourasky Medical Center (16-660-TLV). All patients had a usual interstitial pneumonia (UIP) pattern as determined by expert pathologists (S.B. and E.O.) following international guidelines (1), with multiple fibroblastic foci (FF). Diagnoses were established by multidisciplinary discussion (MDD) in accordance with current guidelines (1-3). Baseline characteristics of these patients can be found in Table E1.

Bronchoalveolar lavage (BAL) was performed on patients with fibrotic interstitial lung disease (ILD) who underwent bronchoscopy as part of their ILD evaluation between 2021-2024. BAL collection, storage and downstream usage was approved by the Institutional Ethics Committee at Tel Aviv Sourasky Medical Center (21-0255-TLV). Informed consent was obtained from all subjects involved in the study. Diagnoses were assigned by MDD following current guidelines (1-3). BAL was carried out in accordance with previous guidelines by the American Thoracic Society (ATS) (4). In summary, the bronchoscope was wedged into a segment of the right middle lobe or lingula. Following that, 150 ml (3x50 ml syringes) of warm (37ºC) sterile saline solution were instilled, with slow aspiration thereafter. The aspirated fluid was collected in a sterile container. Part of the aspirated BAL fluid (~25 ml), which was not required for clinical examination, was processed for research purposes, as described below. Baseline and clinical characteristics of the patients whose BAL fluid was tested for MMP11 levels can be found in Table E2.

**Digital spatial profiling of IPF patient lung tissue**

Formalin-fixed paraffin-embedded (FFPE) lung tissue of an IPF patient was profiled using GeoMx® DSP (5) . Thin (5 µm) tissue sections were prepared and mounted on X-tra positively-charged slides (Leica, Buffalo Grove, IL, USA) according to manufacturer’s recommendations for Semi-Automated RNA Slide Preparation Protocol (FFPE) (manual no. MAN-10151-04). Part of the slides were stained by hematoxylin and eosin (H&E), and were examined by a certified pathologist experienced in ILD (S.B.) to identify regions of fibroblastic foci (FF) and established fibrosis without FF, epithelial, or vascular structures. The consecutive unstained tissue slices were sent to NanoString Technologies laboratories (Seattle, WA, USA) for further processing as further described.

Slides containing lung sections were baked at 65°C prior to going through deparaffinization, rehydration, heat-induced epitope retrieval, and enzymatic digestion. Tissues were then incubated with 10% neutral buffered formalin (NBF) for 5 minutes and 5 minutes with NBF Stop buffer. All steps following baking were carried out on a Leica BOND-RX. Slides were then removed from the Leica BOND-RX and in situ hybridization with GeoMx® Human Whole Transcriptome Atlas (WTA) probes (6), which include probes for 18,677 protein-coding genes and 139 negative control probes, was carried out overnight in a humidified hybridization chamber kept at 37°C. Following rounds of stringent washing with a 1:1 volumetric mixture of 4X Saline Sodium Citrate buffer (4x SSC) and 100% formamide to remove off-target probes, the tissue was blocked with Buffer W blocking solution (NanoString Technologies) then incubated with a cocktail of mouse anti-PanCytokeratin-AF488 (mouse anti-PanCK, Novus NBP2-33200AF488, clone: AE1/AE3), mouse anti-Smooth Muscle Actin-AF594 (mouse anti-SMA, abCam ab202368, clone: 1A4), and mouse anti-CD68-AF647 (mouse anti-CD68, Santa Cruz Biotechnology sc-20060AF647, clone: KP1) antibodies for 60 minutes at RT. The nuclear marker Syto83 was included in the staining cocktail as well.

Tissue morphology was visualized using the fluorescent antibodies and Syto83 on the GeoMx® DSP instrument. Regions of interest (ROIs) were selected from fibroblastic foci (n=4) and established fibrosis (n=9) using the polygon tool. No segmentation of the ROIs was carried out, and UV light was utilized to release and collect oligonucleotides from each ROI. Areas of UV light irradiation within an ROI are referred to as areas of illumination (AOI). During PCR, Illumina i5 and i7 dual-indexing primers were added to each photocleaved oligonucleotide allowing for unique indexing of each AOI. Library concentration was measured using a Qubit fluorometer (Thermo Fisher Scientific), and quality was assessed using a Bioanalyzer (Agilent). The Illumina Novaseq 6000 was used for sequencing, and the resulting FASTQ files were then processed by the NanoString DND pipeline to generate count data for each target probe in every AOI.

**GeoMx DSP analysis**

Initial steps of data filtering and normalization were performed on GeoMx DSP Analysis Suite (V2.3.4). For each ROI, a Limit of Quantitation (LOQ), which served as the background signal, was calculated based on the geometric mean of the signal derived from the negative probes, multiplied by the geometric standard deviation of this signal to the power of 2. Genes that had number of raw reads that was lower than the LOQ were filtered out of the analysis. Then, the data across different ROIs was normalized to the same 3^rd^ quartile (Q3) in the following way. For each ROI, the 75^th^ percentile (Q3) of all probe counts was calculated, and was normalized to the geometric mean across all ROIs to obtain normalization factors for each ROI. Then, raw counts for each gene in each ROI were normalized according to the corresponding normalization factor to obtain normalized counts.

Differential expressed gene analysis on log_2_-transformed normalized counts was performed using linear models by using the R/Bioconductor package limma (7). The design matrix we used was generated by the “model.matrix” function using the ‘~ROI_Name + Slide’ formula. This formula compared between different ROI types, i.e. established fibrosis vs FF, while taking into account the different slides as additional effect. Linear model was then calculated using the “lmFit” function, and p-values were calculated by the empirical Bayes method using the “eBayes” function, which were further adjusted for multiple comparisons by the Benjamini-Hochberg (FDR) correction. FF or fibrosis upregulated genes were considered as genes that exhibit log_2_ Fold-Change (log_2_FC) of at least 0.6 and adjusted p-value (FDR) < 0.05.

**Principal component analysis (PCA)**

PCA on GeoMx spatial transcriptomics data depicted in figure 1B was performed using ‘FactoMiner’ R package (8) ‘PCA’ function with default parameters. PCA visualization was performed using ‘fviz_pca_ind’ function using ‘factoextra’ R package (9).

**Comparison of GeoMx FF-signature to previous IPF spatial transcriptomics datasets**

Our FF-signature, which comprises genes increased in FF compared to established fibrosis, was compared to data reported by Eyres et al. (10) that was also obtained by GeoMx DSP, but using a smaller panel of probes (n=1813). This data included normalized counts of 1813 genes from FF ROIs (n=10) and "adjacent alveolar septae" ROIs (n=10), which served as the closest equivalent of our established fibrosis ROIs, from 3 IPF lung tissues, and can be found in the original manuscript in Table S1. First, we analyzed differentially expressed genes using 'limma' package, similarly to the analysis described in 'GeoMx DSP Analysis' section, except the design matrix accounted for the patient (~ROI_Name + Patient) instead of the slide as in our analysis. In order to account for the smaller probe set that was used by Eyres et al. in their analysis, we only considered genes which were detected in both our probe set and the probe set used by Eyres et al. (n=795) in the calculation of Jaccard similarity and permutation test. Accordingly, FF upregulated genes (FDR < 0.05 and increased in FF, i.e. log_2_FC > 0) in either dataset was only considered if it was part of the 795 shared genes, which yielded 39 genes for Eyres et al. and 48 genes from our FF-signature.

We used a permutation test (number of permutations = 10,000) of the Jaccard similarity between a random sample of 48 genes out of the 795 shared genes and 39 genes out of the shared genes compared to the observed Jaccard similarity for statistical analysis. P-value was reported as p < (b+1) /m, where b corresponds to the number of values that were greater or equal to the observed Jaccard similarity, and m is the total number of permutations.

In addition, we compared our FF-signature to another dataset generated by Kim et al. (11), in which FF ROIs were compared to 'Fibrosis' ROIs using GeoMx and a WTA probe panel. For both Kim et al. FF upregulated genes and our FF-signature we considered all genes that exhibited adjusted p-value (FDR) < 0.05 and were upregulated in FF (log_2_FC > 0), which yielded 274 genes of our FF-signature and 126 genes from Kim et al. dataset. We performed a permutation test (number of permutations = 10,000) between a random sample of 274 genes from the whole set of detected genes in GeoMx (n = 10,185) and a random sample of 126 genes from the whole set of detected genes by Kim et al. (n = 18676) compared to the observed Jaccard similarity. P-value was reported as described above for the other permutation test.

**Comparison of GeoMx FF-signature to FF proteomics data**

Our FF-signature, which comprises genes upregulated in FF compared to established fibrosis was compared to a list of proteins from Herrera et al. (12) that were increased in FF compared to established fibrosis detected by mass spectrometry of regions of interest isolated by laser capture microdissection. As the number of total genes in the proteomic approach that were significantly increased in FF (FDR < 0.05) was 76, we compared them to the top 76 genes in the GeoMx FF-signature, according to log_2_FC. We next used a permutation test (number of permutations = 10,000) of the Jaccard similarity between a random sample of 76 genes of the whole set of detected genes in GeoMx (n = 10,185) and 76 proteins out of the whole set of detected proteins reported by Herrera et al. (n = 2772) compared to the observed Jaccard similarity. P-value was reported as p < (b+1) /m, where b corresponds to the number of values that were greater or equal to the observed Jaccard Similarity, and m is the total number of permutations.

**Re-clustering of scRNA-seq data**

A previous comprehensive dataset by Adams et al. (13), that includes data from 32 IPF and 28 control lungs, was used for deep profiling of FF-fibroblasts that were spatially-resolved using the FF-signature. Data was subset to include only cells that were originally annotated as myofibroblasts, because the FF-signature was detected within this cell population. Unless otherwise stated, throughout the manuscript we refer to these myofibroblasts as fibroblasts. All further downstream processing and analysis was performed using the R package Seurat, v4.0.4 (14). Data was then normalized and scaled to 10000 transcripts per cell using the log-normalization method (15) via the “NormalizeData” Seurat function. The “FindVariableFeatures” function identified the 800 genes with the highest variance to mean ratio using the “vst” selection method. The data was then scaled with the “ScaleData” function for the purpose of principle component (PC) analysis. PCs were calculated using the “RunPCA” function, and the first 20 were further used for the K-Nearest Neighbor (KNN) algorithm using the “FindNeighbors” function. Modularity optimization was performed by the Louvain algorithm using the “FindClusters” function, with resolution parameter set to 0.35. 20 PCs were used for the PHATE dimensionality reduction (16) visualization, with a total of 8 clusters generated using this pipeline. The cluster that encompasses the FF myofibroblasts was identified based on markers that were identified in the GeoMx DSP. Then, differentially expressed genes were analyzed using the “FindMarkers” function, with the minimum fraction of cells (“min.pct” parameter( set to 0.1. *'MMP11+ cluster'* was delineated using Seurat 'CellSelector' tool by selecting MMP11+ cells using a UMAP feature plot depicting *MMP11* expression (see Fig E3C bottom panel). *'MMP11*+ cluster' upregulated genes were considered as genes that exhibit log_2_FC of at least 0.6 and adjusted p-value (FDR) < 0.05.

**Pathway enrichment analyses**

Pathway enrichment analyses for GeoMx genes that were increased in FF and for genes that were upregulated in the ‘*MMP11*+ cluster’ were performed using ‘clusterProfiler’ (17) (version 4.12.6) and ‘ReactomePA’ (version 1.48.0) (18) R packages. The relevant human gene symbols were converted to ENTREZ IDs using ‘bitr’ function using the human gene database ‘org.Hs.eg.db’ (Version 3.19.1). Enriched ‘Reactome’ pathways were computed using the ‘enrichPathway’ function, using the Benjamini-Hochberg p-value adjustment (pAdjustMethod = "BH") for multiple comparisons and p-value cutoff of 0.05. Dot plots were produced using the ‘dotplot’ function.

**RNAscope in-situ hybridization**

In-situ hybridization was performed using RNAscope 2.5 HD Reagent Kit Red (Advanced Cell Diagnostics, Newark, CA, USA) according to manufacturer’s protocol. Five µm-thick sections FFPE tissues were used. Target retrieval was performed at 98-102ºc for 15 minutes and protease treatment was performed for 30 minutes, using manufacturer-supplied reagents and according to manufacturer’s protocol. The Hs-MMP11 (#479741), purchased from Advanced Cell Diagnostics, was used for in-situ hybridization. In addition, positive control probe POLR2A (#310451) and negative control probe targeting bacterial DapB gene (#310043) were used as controls for the in-situ hybridization procedure. Mounting was performed using VectaMount Permanent Mounting Medium (Vector Laboratories) as recommended by the manufacturer. Images were acquired using a Nikon Eclipse Ci microscope (Nikon, Tokyo, Japan) and a Pixelink PL-D755CU camera (Pixelink, Ottawa, ON, Canada), and processed using a Pixelink Capture software (Pixelink).

**Cell culture Transforming Growth Factor β (TGF-β) stimulation**

WI-38 were kindly provided by Dr. Michael Milyavsky (Tel Aviv University, Tel Aviv, Israel). WI-38 were maintained in Minimal Essential Medium (MEM, Biowest, Nuaillé, France), supplemented with 10% fetal bovine serum (FBS, Gibco, Waltham, MA, USA), 1 mM pyruvate (Sigma-Aldrich, St. Louis, MO, USA), 2 mM L-glutamine (Sigma-Aldrich), 100 units/ml of penicillin and 100 µg/ml streptomycin antibiotics (Sigma-Aldrich), and were maintained at 37ºC in a humidified incubator. Cells were checked for the absence of mycoplasma contamination using EZ-PCR Mycoplasma Detection Kit (Sartorius, Göttingen, Germany) according to manufacturer’s instructions. For TGF-β stimulation, cells were seeded in a 6-well plate (300,000 cells/well) in a starvation medium (MEM with no FBS) overnight. Then, cells were treated with 10 ng/ml Human TGF-β1 (Cat. No. 100-21-10, Peprotech, Cranbury, NJ, USA) or vehicle control (PBS) for 24 or 48 hours, after which they were harvested for RNA extraction as described below.

**RNA extraction and reverse transcription quantitative PCR (RT-qPCR)**

Cells were rinsed with PBS and harvested using a cell scraper into an RA1 lysis buffer from the NucleoSpin RNA Mini kit (Cat. No. 740955.50, Macherey-Nagel, Dueren, Germany) or RAL buffer from the Ribospin II (GeneAll Biotechnology, Seoul, South Korea), following an RNA extraction protocol and genomic DNA digestion according to the corresponding manufacturer’s kit instructions. Equal amounts of RNA (300-800 ng) from each sample were used as an input for first-strand cDNA synthesis using the qScript cDNA Synthesis Kit (Cat. No. 95047-100, Quantabio, Beverly, MA, USA) according to manufacturer’s protocol. RT-qPCR was performed on a StepOnePlus real-time PCR System (Thermo Fischer Scientific, Waltham, MA, USA) using Fast SYBR Green Master Mix (Cat. No. 4385612, Thermo Fischer Scientific) and 200 nM of each of reverse and forward primer for each gene. The primer sequences used for RT-qPCR are listed in Table E8. The fold-change calculation between samples was calculated based on a relative standard curve of cDNA sample mix standard using the StepOne Software v2.3 (Thermo Fischer Scientific). Each sample relative quantity of a target gene was normalized to the relative expression of GAPDH housekeeping gene, and the values of each sample were normalized to the corresponding 24h vehicle control-treated sample in each experiment. Statistical significance between vehicle-control treated and TGF-β1 treated sample of each time point was assessed by Student's t-test based on 3 independent experiments.

**MMP11 protein level measurement in bronchoalveolar lavage (BAL) fluid using enzyme-linked immunosorbent assay (ELISA)**

Following acquisition during bronchoscopy, BAL fluid was kept on ice (for up to 1 hour) until it was transferred to the laboratory for processing as further described. The BAL samples were first centrifuged at 300xg for 5 minutes at 4ºC, and the BAL fluid supernatant was aspirated and divided into 2 ml aliquots and stored at -80ºC. On the day of the ELISA experiment, one aliquot of each sample was thawed on ice, then the samples were centrifuged for 20 minutes at 1000xg at 4ºC, and the supernatants were transferred to a fresh tube. MMP11 ELISA was performed on 100 µl of the BAL supernatants using the Human MMP-11 ELISA colorimetric kit (Cat. No. NBP3-06887, Novus Biologicals, Littleton, CO, USA) according to manufacturer’s instructions. The signal was detected as optical density (OD) at 37º and 450 nm wavelength using Tecan Infinite F200 PRO plate reader (Tecan, Männedorf, Switzerland). MMP11 concentration was then calculated based on fitting the OD on a standard curve generated by serial dilutions of an MMP11 reference standard provided by the manufacturer, ranging from 0.16 to 10 ng/ml.

**MMP11 BAL protein levels statistical and survival analyses**

Patients with fibrotic ILD for which MMP11 protein levels in BAL were measured, were divided into a progressive pulmonary fibrosis (PPF) group and a non-PPF group using the 2022 ATS/ERS/JRS/ALAT guidelines definition for PPF (3). In short, progressive disease included at-least two of the following in the year before or after bronchoscopy - 1) worsening respiratory symptoms, 2) absolute decline in FVC ⩾5% predicted or in D_LCO_ (corrected for hemoglobin) ⩾10 %predicted, and 3) radiological evidence of disease progression. Although this PPF definition refers to fibrotic ILD patients other than IPF, we applied it to IPF patients as well for uniformity. The division of patients into PPF and non-PPF group can be found in Table E2. MMP11 levels from patients BAL fluid were presented as median (interquartile range) and compared between patients with and without PPF using the Mann-Whitney U test. The prognostic utility of MMP11 was also evaluated using a time-dependent combined outcome of hospitalization due to ILD exacerbations, lung transplantation and death. Kaplan-Meier and Cox regression analyses were used, with MMP11 protein levels as a continuous variable and as a categorical variable (low/high groups using a cutoff of 1.6 ng/ml). The significance of the difference in survival between the two groups was examined by a log-rank test. All analyses were performed by SPSS software v28 (IBM, Armonk, NY, USA).

**Pseudotime trajectory inference and identification of temporally dynamic genes**

The Seurat object was converted to a ‘SingleCellExperiment’ object using Seurat built-in function ‘as.SingleCellExperiment’. Then, trajectories were inferred using ‘slingshot’ function using the PHATE reduction, and defining that at least one lineage will end at ‘*MMP11*+ cluster’ (end.cluster = ‘FF_cluster’). In order to test for genes that are differentially expressed along the ‘*MMP11*+ cluster’ lineage, we used the ‘tradeSeq’ R package v1.18.0 (19) as described below. For ‘tradeSeq’ analyses, we used only genes that were significantly differentiated expressed (p-value adjusted < 0.05) between the *'MMP11*+ cluster' and the predicted starting cluster (cluster 4) as determined by ‘slingshot’ (number of genes = 1035). In addition, we only analyzed cells that had a weight of at least 0.8 that was assigned to the relevant lineage (n=809). The gene counts from the relevant cells were used to fit a negative binomial generalized additive model (NB-GAM) to the relevant genes as described (19) using the ‘fitGAM’ function and nknots = 5. Out of 1035 genes, 32 did not converge into a fitted model and were excluded from further analysis. The association of gene expression with the pseudotime trajectory was assessed using tradeSeq ‘associationTest’ function, using the following parameters: l2fc = log_2_(1.5) and contrastType = “end”. In addition, we analyzed differentially expressed genes between the beginning and the end of the *'MMP11*+ cluster' trajectory using ‘startVsEndTest’, with l2fc parameter set to log_2_2.

**Transcription factor (TF) analysis**

For TF analysis, we used the union of the genes that were significantly (adjusted p-value < 0.05) associated with the *'MMP11*+ cluster' trajectory as determined by tradeSeq ‘associationTest’ function and the genes that were significantly (adjusted p-value < 0.05) expressed between the beginning and end of the *'MMP11*+ cluster' trajectory as determined by tradeSeq ‘startVsEndTest’ function, a total of 710 genes. In addition, we used only the cells that had a weight of at least 0.8 that was assigned to the '*MMP11*+ cluster' trajectory (n=809). TF analysis was performed using pySCENIC (version 0.12.1) (20) Python (version 3.7.12) package using default parameters as further described. Gene regulatory network, comprising TFs and co-expressed genes that serve as potential target genes, were inferred using grnboost2 algorithm, which were inputted with a count matrix comprising the genes and cells described above. Next, the gene regulatory network modules were pruned based on TF motif enrichment to infer only regulons, which comprise TFs and their direct target genes, using prune2df function. Human Motif2TF annotations v10, Human cisTarget database mc_v10_clust (TSS+/-10kb and 500bpUp100Dw) rankings and hg38 TF list were downloaded from [https://resources.aertslab.org/cistarget/](https://protect.checkpoint.com/v2/r02/___https://resources.aertslab.org/hnxyfwljyd___.YzJlOnRsdm1jOmM6bzpkNzAxOTNiNGNlY2RmZDMxMjRmZWRmZDRkNWExOGNiNDo3OjNhYTg6OWFkMzY5N2ViOTJkODZiOThkOWRiZGNhMzNiZmRlYzk4MWQ1ZDgwNzBhNmUzYWFjNDJmMmQ5ODVjOWUxOTYxNTpwOlQ6VA) and were supplied as reference databases to the aforementioned algorithms. Finally, the regulon activity for each cell was quantified using AUCell function. Regulon activity data was imported to R, the cells were ordered by ‘*MMP11*+ cluster’ lineage pseudotime and heatmaps of AUCell scores were generated using ‘ComplexHeatmap’ R package version 2.20.0 (21).

**TF regulon enrichment and correlation with pseudotime**

TF regulons that were determined by pySCENIC were imported from Python to R using ‘reticulate’ package and ‘read_pickle’ function from the ‘pandas’ (version 1.3.5) Python package. Differential expression between the ‘MMP11 cluster’ and the cluster at the beginning of the predicted trajectory described above (cluster 4) were calculated for all genes using ‘FindMarkers’ Seurat function, and were then ranked in descending order by log_2_FC. Gene-set enrichment analysis was performed by ‘fgsea’ package version 1.30.0 (22) on the differentially expressed genes using the regulons as gene sets that were tested for enrichment. Significantly (p-adjusted < 0.05) enriched regulons were plotted using ‘plotGseaTable’ function. Correlation between TF regulon activation and ‘*MMP11*+ cluster’ lineage pseudotime were calculated using ‘cor.test’ using the Pearson method, and p-values were adjusted for multiple comparisons using Benjamini-Hochberg procedure.

**Cell-Cell Interaction analyses**

The inference of cell-cell interactions that may be relevant for inducing the FF-signature was performed using ‘NicheNet’ algorithm v2.1.0 (23), essentially as described in [nichenetr/vignettes/seurat_steps.md at master · saeyslab/nichenetr · GitHub](https://protect.checkpoint.com/v2/r02/___https://github.com/xfjDxqfgdsnhmjsjywdgqtgdrfxyjwdAnlsjyyjxdxjzwfy_xyjux.ri___.YzJlOnRsdm1jOmM6bzpkNzAxOTNiNGNlY2RmZDMxMjRmZWRmZDRkNWExOGNiNDo3OjRkNzc6ZTE5Y2IzMzFjNzcyODA1Y2M3MTFlZmYxZmE2NzY0OTM0MjIxYzliMmMyNmU4NGQwOGY0ZmYwMTNiYzQ1YzY3OTpwOlQ6VA). Human ligand-target prior model, ligand-receptor network and weighted integrated model were downloaded from [NicheNet-v2: final networks and ligand-target matrices](https://protect.checkpoint.com/v2/r02/___https://zenodo.org/wjhtwixda5a97c6___.YzJlOnRsdm1jOmM6bzpkNzAxOTNiNGNlY2RmZDMxMjRmZWRmZDRkNWExOGNiNDo3OjFhYTY6N2Q3YzJiNWY0ZTQ3NzE5NWJkMThhZDQ3OGY1MTA1YWEzZmY4NWQ2ZDE0ODI0Zjg4NjA5NWJmOWM5NGMzNjMxNzpwOlQ6VA) and were used for analyses. The differentially expressed genes that were used for the analysis were all increased genes between the ‘MMP11 cluster’ and myofibroblast cluster 4 (as described in the TF analysis section) that had an adjusted p-value of 0.05 and log_2_FC of at least 0.5 (n=298). The ligands that were considered were ligands that were expressed in at least 10% of the cells in either the fibroblast, myofibroblast or ABC clusters. The receptors that were considered were receptors that are expressed by 10% of ‘*MMP11* cluster’ cells, and background genes were considered as all genes that were present in the ligand-target matrix that were expressed by at least 10% of ‘MMP11 cluster’ cells (n=7643).

For validation of cell-cell interactions inferred by ‘NicheNet’, we used the ‘CellChat’ v2.1.0 algorithm (24). Overexpressed genes and overexpressed interactions were identified using the ‘identifyOverExpressedGenes’ and ‘identifyOverExpressedInteractions’, respectively. Next, the gene expression data was projected onto protein-protein interaction (PPI) network using the ‘projectData’ function. The probability for interactions was computed using the ‘computeCommunProb’ function, using the ‘truncatedMean’ method, trim parameter set to 0.1, and ‘population.size’ parameter set to ‘FALSE’. In addition, communication probability on the signaling pathway level was computed using ‘computeCommunProbPathway’. Cell-cell interactions involving fewer than 10 cells in either group were filtered out from the analysis. The inferred interactions were plotted using the ‘netVisual_bubble’ function.

**Other computational analyses and visualization**

The heatmap and dendogram of GeoMx data, as well as heatmaps depicting TF activity, were created using ‘ComplexHeatmap’. Visualizations of ‘PHATE’ clusters, including PHATE plots depicting gene expression pattern, were visualized using ‘Seurat’ package functions ‘DimPlot’ and ‘FeaturePlot’. Volcano plots were created using custom code via the ‘ggplot2’ R package v3.5.1 (25). The heatmap shown in Figure E2 was created using ‘Seurat’ package function ‘DoHeatmap’. The Euler and Venn diagrams were plotted using ‘eulerr’ R package v7.0.2 (26). The bar plots depicting RT-qPCR data were plotted using 'ggpubr' package v0.6.0 (27).
